# PAF1C allosterically activates CDK12/13 kinase during RNAPII transcript elongation

**DOI:** 10.1101/2024.10.14.618141

**Authors:** David Lopez Martinez, Izabela Todorovski, Melvin Noe Gonzalez, Charlotte Rusimbi, Daniel Blears, Nessrine Khallou, Zhong Han, A. Barbara Dirac-Svejstrup, Jesper Q. Svejstrup

## Abstract

The mechanisms ensuring temporally correct, site-specific phosphorylation of the RNA polymerase II C-terminal domain (CTD) by cyclin-dependent kinases (CDKs) during the transcription cycle remain poorly understood. Here, we present results from *in vitro* reconstitution of CTD phosphorylation combined with *in vivo* evidence to show that human CDK12 and CDK9 both co-phosphorylate CTD Serine 5 and Serine 2. However, only phosphorylation by CDK12 is stimulated by association with the elongation-specific factor PAF1C, in which the CDC73 subunit contains a short, conserved motif capable of association with and activation of CDK12/Cyclin K. This motif is necessary for cell proliferation and crucial for CTD phosphorylation and transcript elongation. Together, these data provide new insight into basic mechanisms ensuring CDK specificity in the RNAPII transcription cycle.

**One-Sentence Summary:** PAF1C facilitates RNAPII phosphorylation in gene bodies through direct contacts with the active site of CDK12/13.

## Main Text

RNA polymerase II (RNAPII) is responsible for the transcription of all protein-coding genes and numerous non-coding RNAs (*1, 2*). Transcription proceeds in a series of steps, including initiation, promoter-proximal pausing, elongation, and termination, after which RNAPII is recycled for a new round of transcription. During the transcription cycle, different co-factors associate with RNAPII to regulate the progression between steps and to coordinate co-transcriptional events such as mRNA splicing and cleavage/polyadenylation (*3–5*). Co-factor recruitment is governed by phosphorylation of the RNAPII C-terminal domain (CTD), which in humans contains 52 repeats of a heptapeptide comprising Tyr_1_-Ser_2_-Pro_3_-Thr_4_-Ser_5_-Pro_6_-Ser_7_ that is phosphorylated at different sites by the CDKs, CDK7, CDK9 and CDK12/13 and their cognate cyclins. Specifically, transcription initiation is governed by the activity of the CDK7/Cyclin H/MAT1 complex as part of TFIIH, while the transition from promoter-proximal pausing to elongation is triggered by CDK9/Cyclin T1 (also known as pTEFb). Similarly, elongation and termination are regulated by CDK12 and its closely related paralog CDK13 in association with Cyclin K (*6–8*). While CDK7 and CDK9 have been studied for decades, little is known about CDK12 and 13 and their regulation. Importantly, while the target- and site specificity of cell cycle CDKs such as CDK1 and CDK2 is governed by the highly regulated appearance and disappearance of their specific, cognate cyclins through the cell cycle (*9*), the transcription-specific CDKs are constitutively associated with their respective cyclins, yet they all phosphorylate the same main target, namely RNAPII. In light of this, a key question that remains is how the correct RNAPII phosphorylation patterns are achieved during the transcription cycle.

During initiation, TFIIH-associated CDK7 phosphorylates serine 5 (Ser5) of the RNAPII CTD repeats, in a reaction which is highly stimulated by the initiation-specific Mediator complex (*10–12*). This triggers Mediator dissociation (*13*) and the exchange of initiation factors by early elongation/pausing factors (*14*), as well as the recruitment of proteins that specifically recognize the Ser5-phosphorylated (Ser5P) CTD, such as the mRNA capping enzyme (*15, 16*). The subsequent establishment of a promoter-proximal, paused RNAPII complex involves at least two complexes: NELF and DSIF (*17, 18*). CDK9, regulated by a multitude of factors such as the 7SK snRNP complex and AFF4 (*19*), converts this pausing complex into an active RNAPII elongation complex through phosphorylation of the pausing factors and the RNAPII CTD, triggering the release of NELF and the recruitment of elongation factors such as SPT6 and the PAF1 complex (PAF1C) (*20–22*). PAF1C associates with RNAPII and regulates elongation and chromatin structure in a multitude of ways (*23*); it has also been shown to bind CDK9 and CDK12 and to somehow affect promoter-proximal pause release and CTD phosphorylation (*24*). However, the underlying molecular mechanism, and whether these effects are direct, has remained unknown.

Phosphorylation of Ser2 of the RNAPII CTD during elongation has typically been attributed to CDK9 and CDK12/13. Nevertheless, the CTD site-specificity of CDK12/13 and even the identity of a specific “Ser2 kinase” has remained a matter of debate (*25, 26*). Indeed, although phosphorylated Ser2 (Ser2P) represents an abundant mark on the RNAPII CTD in living cells, uncertainty persists about the manner of its formation and the general mechanisms underlying the CTD phosphorylation patterns observed *in vivo*.

In this study, we reconstituted the activity of CDK9/Cyclin T1 and CDK12/Cyclin K complex *in vitro* and show that they predominantly generate Ser2P-Ser5P di-phosphorylated CTDs. Furthermore, we show that CDK12 relies on two main components to achieve full activity, namely its own Arg/Ser (RS)-rich domain and the CDC73 subunit of PAF1C. Indeed, we identify a motif in CDC73 that directly and specifically contacts and activates CDK12/Cyclin K kinase. This new CDC73 motif plays crucial role in maintaining CTD phosphorylation, transcript elongation, and proliferation of human cells.

## Results

### CDK12 and CDK9 generate Ser2 and Ser5 di-phosphorylated RNAPII CTD

Experiments on the CTD site specificity of CDK12/CDK13 have yielded conflicting results depending on the approach and system used (*7, 25, 27–32*). We established an *in vitro* system to study the specificity and mechanism of CDK12/13-mediated phosphorylation using highly purified, recombinant factors, including full-length CDK12 and CDK13/Cyclin K complexes, GST-tagged CTD, and RPB1 from Sf9 (insect) cells. We also isolated RNAPII from pig thymus (fig. S1A). The purified proteins were then used for reconstituted kinase assays, in which the factors were first dephosphorylated, and subsequent *de novo* phosphorylation enabled by the addition of ATP (Fig. 1A). This approach resulted in hyper-phosphorylation of the CTD by both CDK12 and CDK13, as indicated by the slower mobility of the hyper-phosphorylated GST-CTD, RPB1, and RNAPII (Fig. 1B and fig. S1B). As the CDK12 and CDK13 kinases are functionally redundant (*29*), highly similar at the sequence and structural level (*33*), and yielded the same results in all the assays in which we compared them, we focused our subsequent effort on CDK12.

**Fig. 1.**
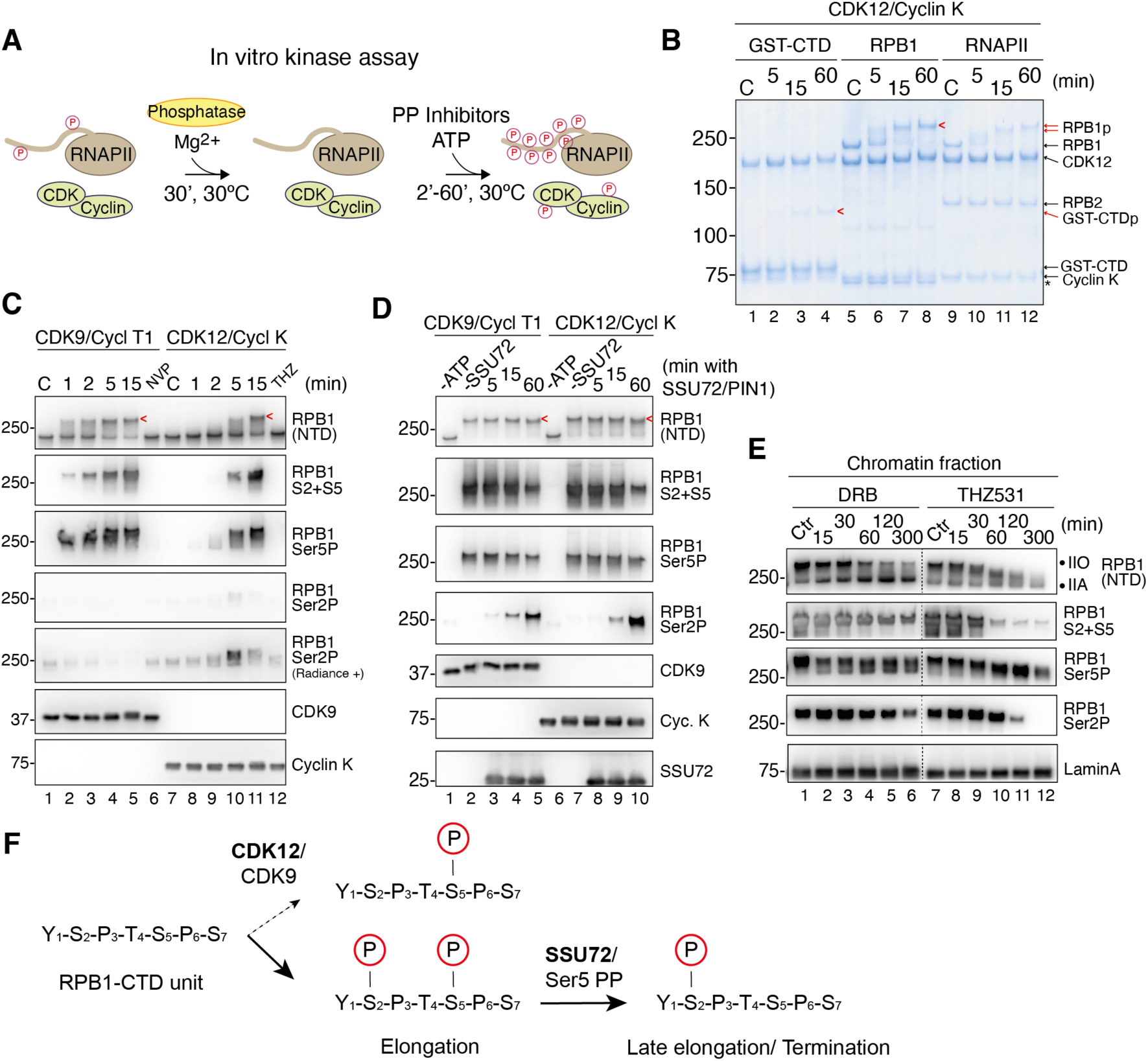
CDK12 and CDK9 generate Ser2 and Ser5 di-phosphorylated RNAPII CTD. (**A**) Schematic representation of the *in vitro* kinase assay. PP: protein phosphatase. (**B**) Coomassie blue-stained SDS-PAGE gel of the components of the *in vitro* kinase assay, incubated for the indicated times in minutes (min). Red arrows indicate phosphorylation products throughout the figures of the paper. C: control with no ATP added. (**C**) Western blot of the *in vitro* kinase assay with CDK9/Cyclin T1 or CDK12/Cyclin K on RNAPII for the indicated times. Ser5P (3E8 antibody), Ser2P (3E10), and Ser2P+Ser5P (S2+S5) (D1G3K) were examined. CDK-specific inhibitors were added for validation (NVP-2 for CDK9; THZ531 for CDK12). C: no ATP; NVP: 2,5 μM NVP-2 added; THZ: 2,5 μM THZ531 added. NTD: N-terminal domain. (**D**) Western blot of the *in vitro* kinase assay with CDK9/Cyclin T1 or CDK12/Cyclin K on RNAPII followed by dephosphorylation by SSU72/PIN1 for the indicated times. -ATP: no ATP; -SSU72: no SSU72 added. (**E**) Western blot of MRC5 chromatin fractions treated with DRB (100 μM) or THZ531 (1 μM) for the indicated times. Ctr: no drug added. (**F**) Schematic representation of the proposed mechanism for the generation of Ser2P CTD during late elongation. Following Ser2+Ser5 di-phosphorylation by CDK12 or CDK9, Ser5P can be dephosphorylated by SSU72 or other Ser5-specific protein phosphatases (Ser5 PP).

Previous studies on CDK12/13 were typically performed with proteins missing the N- and C-terminal intrinsically disordered regions (IDRs) (*26, 34*). We generated truncated forms that miss one or more of these domains (fig. S1C). Upon expression and purification of CDK12 and its sub-forms in complex with Cyclin K (fig. S1D), *in vitro* kinase assays revealed that only full-length (WT) CDK12 or a version lacking the C-terminal IDR were distinctly active, with the N-terminal IDR and in particular the arginine-serine rich (RS) domain necessary for full CDK12 activity (fig. S1E). One possibility is that the RS domain helps establish contacts with the RPB1 CTD. To address this possibility, we investigated CDK12/Cyclin K binding to immobilized GST-CTD (fig. S1F). Only the full-length CDK12 protein showed good retention on beads. Moreover, in the absence of the RS domain, CDK12 was washed off the immobilized GST-CTD beads, while WT CDK12 remained bound, even at higher NaCl concentrations (fig. S1F). These data open the possibility that the RS domain promotes CDK12 activity at least partly by stabilizing binding to the CTD.

The site-specificity of CDK12/Cyclin K was now compared with that of purified CDK9/Cyclin T1 (fig. S2A). Surprisingly, while rapid and robust formation of Ser5P was observed with CDK9 and somewhat less with CDK12, very little Ser2P was generated (Fig. 1C, lane 5 and 10). Given that CDK9 and CDK12 are generally considered Ser2 kinases, this result was surprising. Intriguingly, however, a marked reduction of the already low level of Ser2P was observed at the last timepoint in the CDK12 assay (Fig. 1C, compare lane 10 with lane 11), leading us to speculate that antibody recognition of the Ser2P mark might be obstructed by Ser5P addition. We therefore checked CTD phosphorylation with an antibody directed against <u>di</u>-phosphorylated Ser2 and Ser5 marks (Ser2P-Ser5P). A strong signal for Ser2P-Ser5P phosphorylation was indeed observed with both CDK12 and CDK9 (Fig. 1C, RPB1 S2+S5). While CDK9 appears to first phosphorylate Ser5, with Ser5-Ser2 di-phosphorylation appearing slightly later, CDK12 appears to generate Ser5P and Ser2P in a highly coupled reaction, with the occurrence of Ser2P-Ser5P di-phosphorylation mirroring that of Ser5P, and with very little Ser2 signal detected (Fig. 1C, lanes 8-11). Phosphorylation of Ser2 by CDK9 and CDK12/13 thus appears to only rarely occur on its own but is instead tightly coupled to Ser5 phosphorylation. ELISA assays with chemically synthesized CTD peptides corroborated that the frequently used Ser2P antibody 3E10 cannot recognize Ser2P when it is part of a Ser2P-Ser5P di-phosphorylated CTD (*35*), and also indicated the specificities of the 3E8 (Ser5P) and D1G3K (Ser2P-Ser5P) antibodies (fig. S2B-C).

To further investigate whether Ser2P marks are generated by CDK9 and CDK12 but obstructed from antibody detection by the near-simultaneous Ser5 phosphorylation, we performed kinase assays followed by Ser5P de-phosphorylation by the Ser5-specific CTD phosphatase SSU72 (stimulated by the prolyl isomerase PIN1 (*36*); see fig. S2A, lanes 2 and 3). SSU72-mediated dephosphorylation indeed resulted in a reduction of the Ser2P-Ser5P signal, and to a smaller extent Ser5P, parallelled by a strong increase in the Ser2P signal (Fig. 1D lanes 5 and 10, respectively). Taken together, these results indicate that CDK12 and CDK9 co-phosphorylate Ser5 and Ser2, and that CTD Ser2 mono-phosphorylation only becomes easily detectable upon Ser5P de-phosphorylation. This illustrates the challenge of correctly interpreting CTD phosphorylation signatures, with many of these signatures over the years having been generated with the phosphorylation-specific antibodies used here (*35, 37*).

We now addressed the question of whether a similar mechanism might be in play *in vivo*, mindful that previous experiments on the effect on CTD phosphorylation of CDK12/13 loss yielded conflicting results (*7, 25, 27–32*). For this purpose, a time-course inhibition of CDK9 (using the inhibitor DRB) and CDK12/CDK13 (using THZ531) was performed, during which the occurrence in chromatin of different RNAPII-CTD phospho-marks was investigated (Fig. 1E). While both inhibitor treatments led to a reduction in Ser2-Ser5 di-phosphorylation, CDK12/CDK13 inhibition led to a faster and much more complete loss of the mark. The partial loss of Ser2P-Ser5P in DRB-treated cells does not appear to be caused by incomplete CDK9 inhibition since the more potent CDK9 inhibitor NVP-2 showed a similar pattern (Fig. S2D) but might instead suggest a stronger dependence of global Ser2P-Ser5P levels on CDK12/13 activity. Intriguingly, the effects of CDK12/13 inhibition on CTD hyper-phosphorylation, i.e. gradual loss of the IIO form, was accompanied by a temporal increase in Ser5P, which eventually decreased to very low levels. Likewise, a temporal increase in Ser2P preceded a clear reduction in the same signal, but this reduction was observed much later than for Ser2P-Ser5P (Fig. 1E, lanes 7-12),. These results illustrate how, at steady-state, CTD phosphorylation levels at different serine residues are reached by a complex interplay between CTD kinases and phosphatases. More importantly, they support the idea that Ser2P-Ser5P di-phosphorylated RNAPII-CTD is an important and surprisingly abundant mark in cells, and that Ser2P may frequently derive from the Ser2P-Ser5P mark through Ser5P dephosphorylation (Fig. 1F).

### The CDC73 subunit of PAF1C specifically activates CDK12 through a direct interaction

Having observed that both CDK9 and CDK12 co-phosphorylate CTD Ser5 and Ser2, we now sought to address how these CDKs exert their function in a temporally correct and potentially distinct manner. One functional discriminator would be the effect of co-factors. Previous data had demonstrated an interaction of elongation factor PAF1C with CDK12 and CDK9, respectively (*24*). We confirmed the CDK12-PAF1C interaction by performing immunoprecipitation (IP) of Flag-tagged CDK12 from HEK293 cell extracts, which co-precipitated PAF1C subunits (Fig. 2A). To evaluate whether this interaction is direct, we expressed and purified the five-subunit PAF1C (Fig. 2B; see fig. S3A, lane 1) and performed a CDK12 binding assay. PAF1C (St-CTR9) co-IPed CDK12 independently of CDK12’s IDRs, indicating that the interaction is direct and occurs through the CDK12 kinase domain and/or Cyclin K (Fig. 2C).

**Fig. 2.**
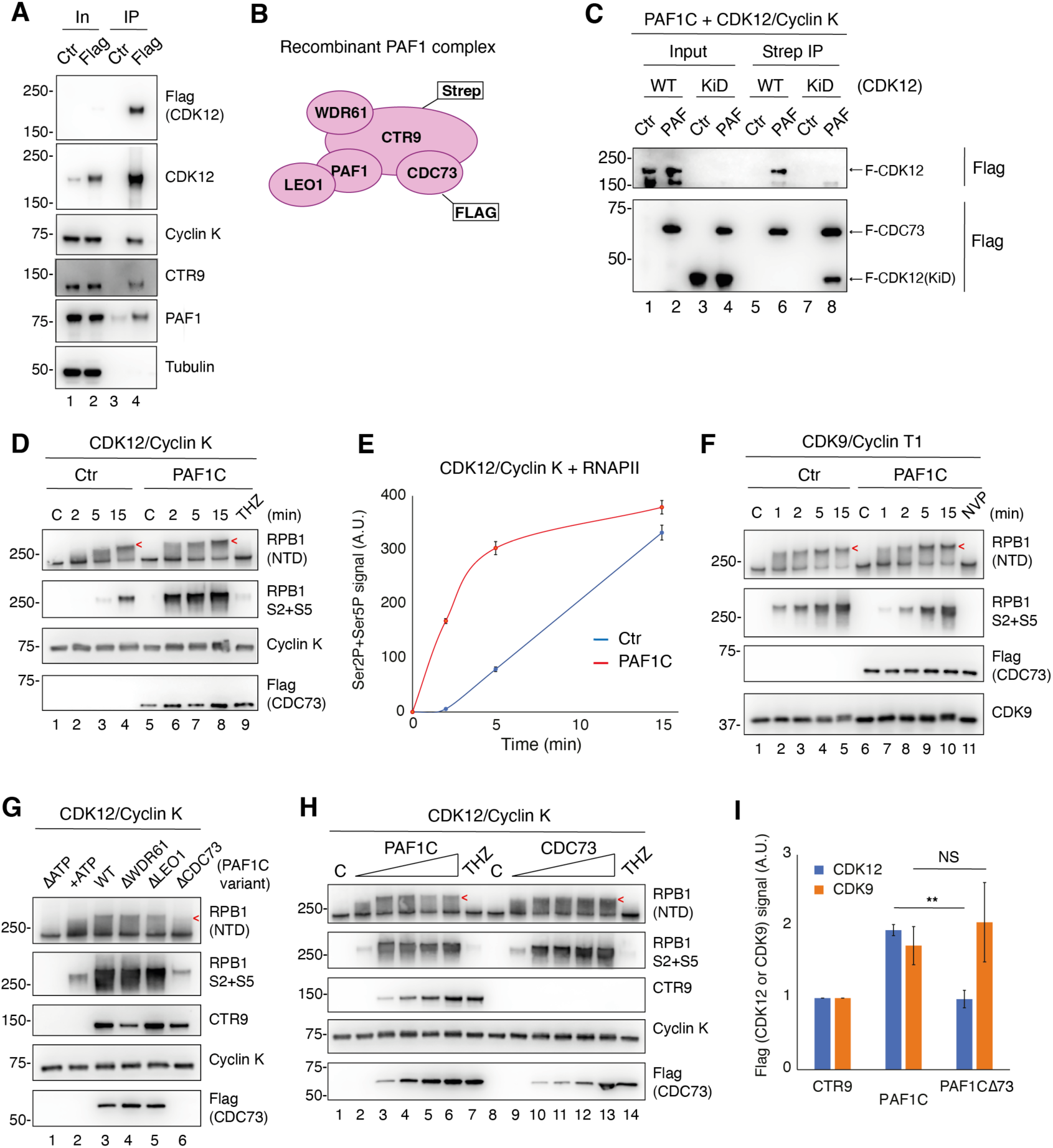
PAF1C subunit CDC73 directly contacts and specifically activates CDK12. (**A**) Western blot of the Flag IP from HEK293 cells expressing Flag-HA-CDK12. In: Input, Ctr: no Flag-HA-CDK12 expressed, Flag: Flag-HA-CDK12 expressed. (**B**) Schematic of PAF1C and tags used for purification from insect cells. (**C**) Western blot of the *in vitro* binding reaction, mixing PAF1C and the indicated forms of CDK12/Cyclin K, followed by Strep IP. Ctr: no PAF1C added. (**D**) Western blot of an *in vitro* kinase assay using CDK12/Cyclin K and RNAPII complemented with PAF1C for the indicated times. Ctr: no PAF1C added, THZ: 2,5 μM THZ531 added. NTD: N-terminal domain, S2+S5: Ser2P + Ser5P. (**E**) Quantification of an *in vitro* kinase assay using CDK12/Cyclin K and RNAPII comparing the presence and absence of PAF1C in the reaction in (D) (n = 3 independent experiments). Signal is mean with error bars indicating standard error. A.U: arbitrary units. (**F**) Western blot of an *in vitro* kinase assay using CDK9/Cyclin T1 and RNAPII complemented with PAF1C for the indicated times. As in (D). (**G**) Western blot of an *in vitro* kinase assay using CDK12/Cyclin K and RNAPII complemented with PAF1C WT or incomplete variants for 5 minutes. ΔATP: no ATP. (**H**) Western blot of an *in vitro* kinase assay using CDK12/Cyclin K and RNAPII complemented with PAF1C or Flag-CDC73 titrated in for 5 minutes. C: no ATP added, THZ: 2,5 μM THZ531 added. (**I**) Quantification of an *in vitro* binding assay as in (C). Mean signal from CDK12 or CDK9 (Flag) was normalized to St-CTR9 (negative control) to show the specific binding (n = 3 independent experiments). Signal is mean, with errors bars indicating standard error. NS: not significant and **: p < 0,01 as determined using an unpaired t-test.

To assess a possible functional role of this interaction, kinase assays were performed using purified RNAPII and CDK12/Cyclin K, in the absence or presence of PAF1C (Fig. 2D). Interestingly, the addition of PAF1C dramatically stimulated the reaction (Fig. 2D, compare lanes 6-8 with lanes 2-4; quantification in Fig. 2E). Since PAF1C has been shown to interact with CDK9 as well (*24*), we also performed a binding assay with PAF1C and CDK9/Cyclin T1 and confirmed their direct interaction (fig. S3B). Strikingly, however, we consistently observed no stimulatory effect of PAF1C on CDK9 kinase activity (Fig. 2F). Together, these data indicate that PAF1C directly and specifically stimulates CDK12 activity *in vitro*.

To investigate which PAF1C subunit is responsible for CDK12/Cyclin K stimulation, we expressed and purified PAF1Cs missing one different peripheral subunit at a time: WDR61, LEO1 or CDC73 (see fig. S3A, lanes 2-4). We then performed CDK12 kinase assays complemented with WT or the various incomplete PAF1Cs. Only PAF1C missing the CDC73 subunit failed to stimulate the reaction (Fig. 2G), indicating that this subunit is required for CDK12 kinase activation. To test if CDC73 is also sufficient to stimulate CDK12 activity, we complemented the kinase reaction with increasing amounts of PAF1C or CDC73 alone, respectively. Remarkably, stimulation of Ser2P-Ser5P phosphorylation was observed with both (Fig. 2H, compare lanes 2 and 9 with lanes 3-6 and 10-13, respectively). These data indicate that the CDC73 subunit of PAF1C is thus both necessary and sufficient to activate CDK12.

To evaluate if CDC73 is also responsible for mediating direct interactions between CDK12/Cyclin K and PAF1C, we performed *in vitro* binding assays. CDK9/Cyclin T1 and the PAF1C scaffold subunit CTR9 were included as a positive and negative controls, respectively. In the absence of CDC73, the interaction between CDK12/Cyclin K and PAF1C was greatly reduced (Fig. 2I and fig. S3C). On the other hand, the interaction between CDK9/Cyclin T and PAF1C was unaffected by the absence of CDC73 (Fig. 2I and fig. S3D). Together, these results show that CDC73 specifically and directly interacts with CDK12 to stimulate its kinase activity, potentially in an allosteric manner.

### CDC73 contains a specific motif that interacts with Cyclin K and CDK12 and promotes the opening of the CDK12 T-loop

We now employed AlphaFold-Multimer (AF2) (*38–40*) for *in silico* protein folding predictions of CDK12 KiD, Cyclin K, and CDC73. This yielded a high confidence structural model of a trimer (Fig. 3A and fig. S4A), where the region of CDC73 predicted to contact CDK12 KiD and Cyclin K maps to a short, previously unstudied motif between residues 290 and 324 (fig. S4B-C). This CDC73 motif expands through a groove in Cyclin K and ends in the proximity of the CDK12 kinase domain (Fig. 3A). As such, we named it the Cyclin K-interacting motif (KIM). Further examination of the predicted structure revealed that the N-terminal portion of the CDC73-KIM is in direct contact with the activation T-loop of CDK12 in its open, or active, conformation, with threonine 893 (T893) in the CDK12 T-loop directly stacked against CDC73 Y290 and Y293 (Fig. 3A, zoomed-in panel). This suggests that the CDC73-KIM elicits an open T-loop conformation and points to a structural basis for allosteric activation of CDK12 by CDC73.

**Fig. 3.**
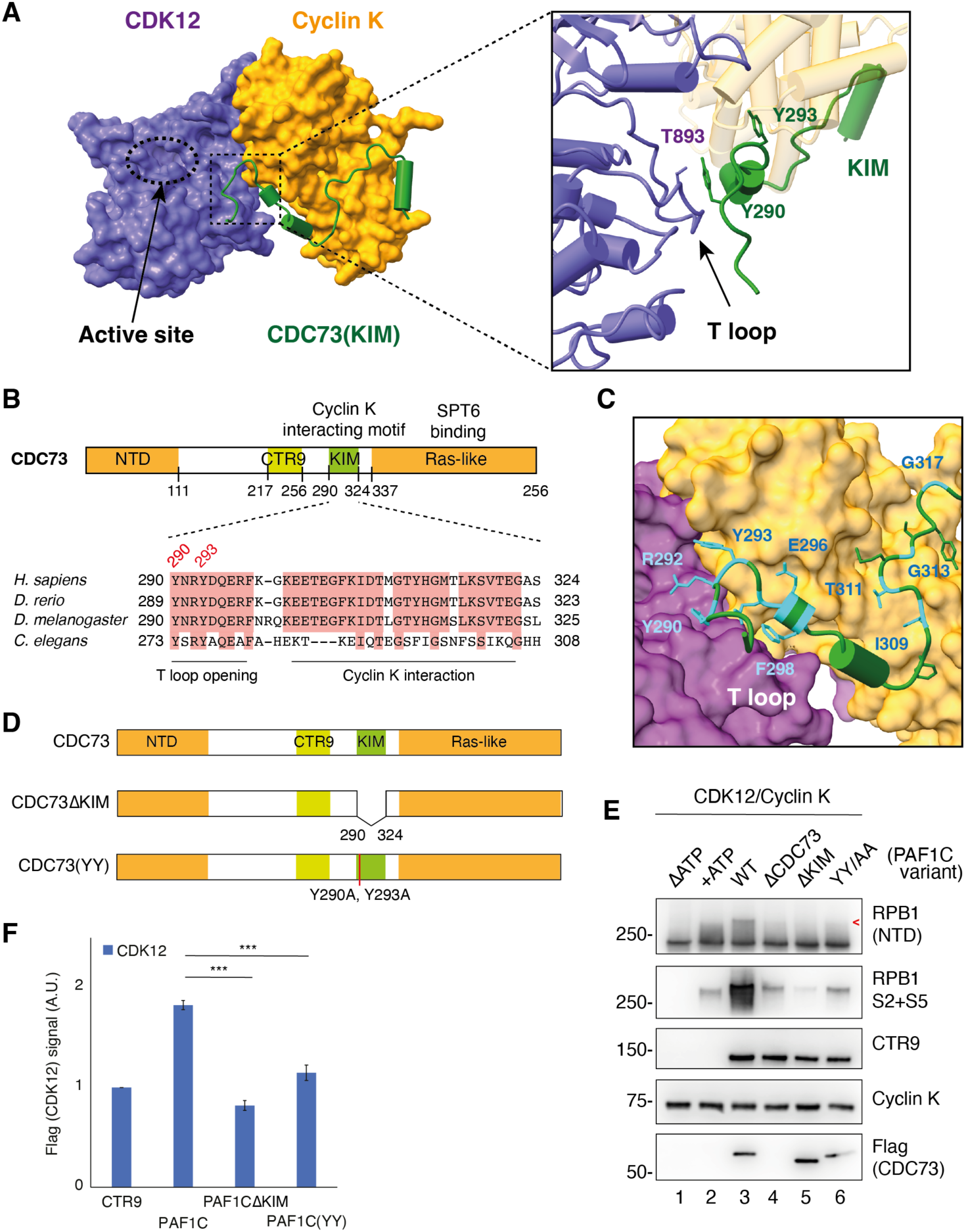
CDC73-KIM interacts with both Cyclin K and CDK12 and promotes the opening of the CDK12 T-loop. (**A**) (Left) AlphaFold2 structural model of CDK12 kinase domain (purple), Cyclin K cyclin fold (yellow) and CDC73-Cyclin K interacting motif (KIM; green). The CDK12 active site indicated. (Right) Zoom-in showing T893 in the CDK12 T-loop in direct contact with Y290 of the CDC73-KIM. (**B**) (Top) Schematic of CDC73 domain organization. NTD: N-terminal domain. (Bottom) Multiple sequence alignment of CDC73-KIM in humans (*Homo sapiens*), zebrafish (*Danio rerio*), fruit fly (*Drosophila melanogaster*) and the worm (*Caenorhabditis elegans*). Fully conserved residues shaded in red. KIM regions important for T-loop opening and the Cyclin K interaction indicated. (**C**) Zoom-in on AF2 structural model of CDK12 (purple)/Cyclin K (yellow)/CDC73-KIM (green and light blue) complex from (A), highlighting in light blue the fully conserved residues as determined in (B). (**D**) Schematic of the CDC73 mutants: CDC73ΔKIM (missing residues 290-324) and CDC73-YY/AA (Y290A, Y293A). (**E**) Western blot of an *in vitro* kinase assay using CDK12/Cyclin K and RNAPII complemented with PAF1C WT, CDC73 absent (ΔCDC73) or the CDC73 mutant variants shown in (D). ΔATP: no ATP added. NTD: N-terminal domain, S2+S5: Ser2P + Ser5P. (**F**) Quantification of the *in vitro* binding assay using Strep-CTR9, PAF1C, PAF1CΔKIM or PAF1C(YY) and the CDK12(KiD)/Cyclin K complex. Mean signal from CDK12 (Flag) was normalized to St-CTR9 (negative control) to show the specific binding (n = 3 independent experiments). Signal is mean with error bars indicating standard error. NS: not significant and ***: p < 0,001 as determined using an unpaired t-test.

Multiple sequence alignment across four distant metazoan species revealed that the CDC73-KIM is highly conserved, with Y290 and Y293 fully conserved to *C. elegans* (Fig. 3B). In general, the highly conserved residues (in light blue in Fig. 3C) perfectly match the amino acids that provide key interactions in the model (such as E296 and F298), or they correspond to small glycine residues (G313 and G317) that provide flexibility to fit in the Cyclin K groove (Fig. 3C). The CDC73-KIM is not conserved in yeast, but the CDK12 ortholog in *Saccharomyces cerevisiae*, Ctk1, is part of a trimer with Ctk2 and Ctk3. Intriguingly, Ctk3 (which is not found in humans) was previously shown to directly open the T-loop of the Ctk1 kinase in a manner similar to that described above for the CDC73-KIM and CDK12 (*41*) (fig. S4D). This may constitute an interesting case of parallel evolution, and potentially has important implications for different modes of regulation employed during transcription elongation in lower eukaryotes and metazoans, respectively.

To further explore the functional significance of the CDC73-KIM in the broader context of transcription, we overlayed the predicted CDC73, CDK12 and Cyclin K trimer onto the previously resolved human RNAPII elongating complex (*42*). Gratifyingly, the position of the CDC73-KIM between the CTR9 interaction and the SPT6 binding domains in the C-terminus of CDC73 (*43, 44*) (Fig. 3B, upper), enables the placement of CDK12/Cyclin K in close proximity to the RNAPII CTD, ready for action (see fig. S4E). This model further supports the notion that CDK12 is allosterically stimulated to phosphorylate the CTD specifically when RNAPII is associated with PAF1C during active elongation.

To validate the AlphaFold model, we first expressed and purified PAF1Cs containing different CDC73 variants, namely WT, CDC73 lacking the KIM (CDC73ΔKIM), or CDC73 with alanine mutations at both location Y290 and Y293 (CDC73-YY/AA) (Fig. 3D and fig. S4F). These proteins were then used to assess the effect on CDK12 kinase activity. Remarkably, a complete loss of stimulatory effect on kinase activity was observed with CDC73ΔKIM- and CDC73-YY/AA-containing PAF1Cs, as determined by the lack of low-mobility RPB1 isoforms and reduced Ser2P-Ser5P signal (Fig. 3E). As the AlphaFold model indicates a direct CDC73-KIM interaction with CDK12/Cyclin K, we also assessed the effect on this interaction. Deletion of the CDC73-KIM caused a loss of the PAF1-CDK12/Cyclin K interaction similar to that observed without the CDC73 subunit, while the CDC73-YY/AA mutant showed a partial defect (Fig. 3F). These results are consistent with a model in which the CDC73-KIM is required for both the interaction with, and the allosteric activation of, CDK12 by PAF1C *in vitro*.

### CDC73-KIM is crucial for CTD phosphorylation, transcript elongation and cell proliferation

Given the key role of CDK12 and CDK13 activity for gene expression, particularly in long genes (*28, 34, 45*), we investigated the role of the CDC73-KIM in cells. For this purpose, we first established a ‘switchover’ system where endogenous CDC73 is knocked down by siRNA in Flp-In T-Rex HEK293 cells, and different recombinant, siRNA resistant, and exogenous Flag-tagged versions of CDC73 are induced by doxycycline (Dox) addition (Fig. 4A and fig. S5A-B). Using this system, we first measured cell growth over time. As expected, a loss of cell proliferation was observed upon CDC73 depletion (Fig. 4B). Importantly, while the phenotypic effects of CDC73 depletion could be rescued by CDC73-WT expression, this was not the case with neither CDC73ΔKIM nor CDC73-YY/AA (Fig. 4B, and fig. S5C), suggesting that the interaction between PAF1C and CDK12/13 is essential for cell proliferation.

**Fig. 4.**
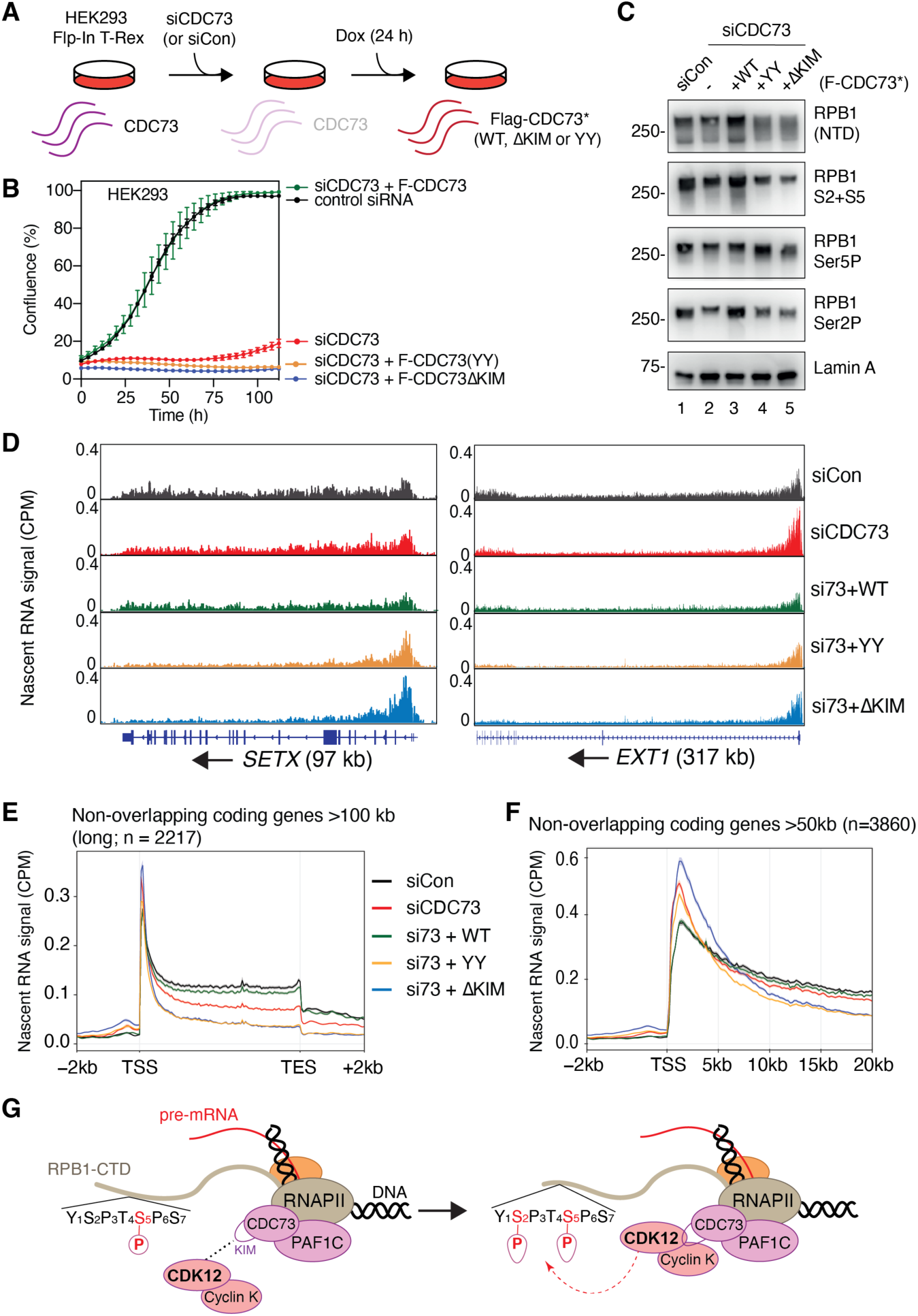
CDC73-KIM is crucial for CTD phosphorylation, transcript elongation and cell proliferation. (**A**) Schematic of the switchover system. Following knock-down of endogenous CDC73, expression of the recombinant, siRNA resistant, Flag tagged CDC73 (Flag-CDC73*) is induced with doxycycline (Dox; 1µg/mL) for 24 hours (h). siCDC73: siRNA targeting CDC73. siCon: siRNA targeting control locus. (**B**) Cell confluence (in (%)) measured over time using the switchover system. Cell confluence was tracked every 4 h for 5 days. Point indicates mean of technical duplicate and error bars indicate standard deviation. Image is representative of experiment performed in biological triplicate. F: Flag. (**C**) Western blot of chromatin fractions following switchover, measuring total RPB1, Ser2P, Ser5P and Ser2P-Ser5P signals. (**D**) Representative images of nascent RNA read counts across *SETX* and *EXT1*, with the direction of transcription indicated by arrow. (**E**) Metagene analysis of non-overlapping coding genes longer than 100 kb (n = 2217) using library- and spike-in normalized nascent RNA read counts obtained from transient transcriptome chem sequencing (TTchem-seq). Line indicates mean signal of experiment performed in technical triplicate. Shaded area indicates standard error. Si73: siRNA targeting CDC73, CPM: counts per million, TSS: transcription start site, TES: transcription end site, kb: kilobases. (**F**) Same as (E) except metagene analysis was performed using non-overlapping coding gene intervals longer than 50 kb (n = 3860) showing the TSS and first 20 kb. (**G**) Schematic of PAF1C-mediated CDK12 activation during transcript elongation.

We also investigated the potential role of the CDC73-KIM in CTD phosphorylation (Fig. 4C). Interestingly, expression of CDC73ΔKIM or CDC73-YY/AA led to a clear reduction in the levels of Ser2P and Ser2P-Ser5P di-phosphorylated CTD (Fig. 4C, compare lanes 4 and 5 with lane 3), to an extent which is highly reminiscent of the CTD phosphorylation defects observed upon chemical inhibition of CDK12 and CDK13 in cells (*cf.* Fig. 1E). These data indicate that CDC73-KIM is indeed important for the activation of CDK12/13 also *in vivo*. As a defect in CTD phosphorylation would be expected to impact transcription, we next examined the role of the CDC73-KIM in nascent RNA production using transient transcriptome sequencing, TT_Chem_-seq (*46, 47*). While transcription remained robust in the beginning of genes, a relative loss of activity was observed across the gene body and towards the 3’-end, including in the *SETX* and *EXT1* genes, upon CDC73 depletion, but dramatically more so upon expression of the CDC73-KIM mutants (Fig. 4D). Metagene analysis showed that this was a genome-wide effect, most clearly observed at long genes (>100 kilobases; kb; Fig. 4E) but generally across different gene groups (fig. S5D), except for short genes (<10 kb), which were less affected (fig. S5E, left panel), or sometimes even upregulated, such as *GADD45B* and *HSPA1A* (fig. S5E, right panels). A more detailed metagene analysis of the first 20 kb of long genes demonstrated that, while initiation and CDK9-dependent 5’-end transcription appeared to be largely unaffected or potentially even relatively increased, a distinct drop in the nascent RNA signal 5-10 kb into the gene body in genes occurred in cells expressing CDC73ΔKIM or CDC73-YY/AA, (Fig. 4F). Together, these data point to a crucial role for the CDC73-KIM for transcript elongation and thus transcription of long genes in human cells.

## Discussion

While Ser2P-Ser5P di-phosphorylated CTD has previously been detected in cells (*48*), this mark has been largely disregarded in most studies of RNAPII CTD phosphorylation in humans. Our data open the possibility that the Ser2P-Ser5P mark is both highly abundant and dependent on CDK12/13 and CDK9 activity. We propose that the biological significance is twofold. First, Ser2P-Ser5P CTD may allow specific interaction of certain CTD-binding domains such as SRI (Set2 Rpb1-Interacting) domains, present in proteins like SETD2 (*49*), or some CIDs (CTD-interacting domains), such as those in SCAF4 and SCAF8 (*50*), but also that it acts as a precursor for the Ser2P CTD mark, which we suggest may be generated primarily through Ser5P de-phosphorylation towards the end of genes. In fact, the Ser5P-specific phosphatase SSU72 is part of the 3’-cleavage machinery recruited to RNAPII late in the transcription cycle, which might help explain how, somewhat paradoxically, SSU72 generates elevated Ser2P levels in this region (*51*), and also how Ser2P and 3’-processing are mechanistically coupled in human cells (*52*).

Another interesting question is whether and how CDK9 or CDK12 cooperate to generate and maintain CTD marks in cells. We favor a model in which CDK9 acts as the *de novo* Ser5P-Ser2P kinase, while CDK12/13 is required for maintaining and expanding this CTD mark during transcription elongation, particularly across the long genes of human cells. In budding yeast, Ctk1 is the likely functional homologue of CDK12/13. Interestingly, while Ctk1 is dispensable for yeast viability (*53*), CDK12/13 are essential in human cells (combined loss is lethal (*29*)). We suggest that this difference may reflect that yeast genes are short (1,5 kb long on average (*54*)) so that Ctk1 activity is not absolutely required, while CDK12/13 are essential to enable processive transcript elongation across the much longer genes in humans (24 kb on average (*55*)).

We show that CDK12/13 is bound and activated by PAF1C through a short, conserved motif found in the CDC73 subunit. This in turn is necessary for global transcription, particularly for long genes which have an increased dependence on elongation for their efficient transcription, a role that therefore cannot be compensated for by CDK9. Interestingly, several reports have shown that RNAPII transcript elongation rates vary significantly within individual genes, starting as low as 0.5 kb/min in the first ∼10 kb, but accelerating to 2–4 kb/min further downstream (e.g. (*55–58*). We note that this region of fast elongation coincides remarkably well with the region of genes requiring the CDC73 KIM for efficient transcription. This supports a working model in which the establishment of PAF1C-mediated stimulation of CDK12/13 activity helps accelerate RNAPII elongation.

Our characterization of the CDC73 KIM suggests a mechanism for activation of a transcriptional CDK that is dynamic, and specifically coupled to transcript elongation. Other CDK activation mechanisms include the previously described Ctk3-Ctk1 complex in yeast (*41*), but also CDK7 kinase, where direct contacts between the CDK7 T-loop and MAT1 were shown to promote CTD phosphorylation (*59, 60*). However, in these examples, the ‘activators’ are constitutively bound to the relevant CDK/Cyclin complex and therefore not subject to the temporal and spatial regulation that the CDC73-KIM exerts on CDK12. Intriguingly, we note that activation of CDK9 by the HIV-1 protein Tat (*61*) is somewhat similar to the CDK12/CDC73-KIM described here. For viral CDK9 activation, the Tat protein interacts with Cyclin T1 and extends onto the T-loop of CDK9, stacking on top of it and stabilizing the open conformation of the kinase, resulting in stimulation of CTD phosphorylation (*62*).

In conclusion, we have uncovered a novel mechanism for the elongation-specific activation of the CTD kinase CDK12 via the CDC73 subunit of PAF1C, in which a previously unstudied motif in CDC73 enables the allosteric stimulation of CDK12 for Ser2P-Ser5P di-phosphorylation (Fig. 4G). The *in vitro* reconstitution and *in vivo* switchover system described here pave the way to better understand the division of labor between the different transcriptional CDKs, characterize their specific substrates, and help further unravel molecular mechanisms of transcription.

## Acknowledgments

We thank Dhira Joshi and Nicola O’Reilly at The Francis Crick Institute for CTD peptide synthesis. We also thank all the members of the Svejstrup lab and CGEN for insightful discussions.

## Funding

Novo Nordisk Foundation grant NNF19OC0055875 (JQS)

Danish National Research Foundation Chair grant DNRF153 (JQS)

Danish National Research Foundation Center of Excellence grant DNRF166 (JQS)

EMBO Postdoctoral Fellowship ALTF 847-2023 (IT)

EMBO Postdoctoral Fellowship ALTF 2020-260 (MNG)

European Union’s Horizon 2020 Marie Sklodowska-Curie grant 101023584 (MNG)

## Author contributions

Conceptualization: DLM, JQS

Methodology: DLM, IT, MNG, CR, DB, NK, ZH

Investigation: DLM, IT, MNG, CR, DB, NK, ZH

Visualization: DLM, IT, MNG

Funding acquisition: JQS

Project administration: DLM, JQS

Supervision: JQS

Writing – original draft: DLM, JQS

Writing – review & editing: DLM, ABDS, JQS

## Competing interests

Authors declare that they have no competing interests.

## Data and materials availability

Genome-wide data that support the findings of this study have been deposited in GEO (https://www.ncbi.nlm.nih.gov/geo/) with the accession codes GSEXXXXX. All other data are available in the main text or the supplementary materials.

## Supplementary Materials

### Materials and Methods

#### Cloning and protein expression

Protein and complex expression was performed in Sf9 insect cells following the biGBac method (*1, 2*). Full-length, human cDNAs were cloned from RNA extracted from RPE1 or U2OS cells and subcloned into pLIB plasmids. Primers included the tag sequence used and a 3C Prescission protease target site. Factors used in this study and their tag (position in parenthesis) included: Flag(N)-CDK12, Flag(N)-CDK13, Cyclin K, Flag(N)-CDK9, Cyclin T1, Flag-GST(N)-RPB1-CTD(1586-1971), Flag(N)-RPB1, His(N)-SSU72, Strep(II)(N)-PIN1, Strep(II)(C)-CTR9, Flag(N)-CDC73, LEO1, PAF1, WDR61. Individual cDNAs were combined into pBIG1 plasmids according to their complexes: CDK12/Cyclin K, CDK13/Cyclin K, CDK9/Cyclin T1, PAF1 complex. pLIB containing full-length versions of the cDNAs were used as templates to generate truncated or mutated forms: CDK12ΔNTD (700-1490), CDK12ΔCTD (1-1082), CDK12-KiD (700-1082), CDK12ΔRS (Δ104-380), CDC73ΔKIM (Δ290-324) and CDC73-YY/AA (Y290A, Y293A).

pLIB or pBIG1 plasmids were used for bacmid generation in DH10BAC. Bacmids were transfected into Sf9 cells with ExpiFectamine (Gibco) according to manufacturer’s instructions and baculovirus (P1) were harvested from the supernatant after 5 days. Sf9 cells were infected with between 0.1 – 0.002% P1 baculovirus and supernatant harvested after 4 days as P2 baculovirus. 1 % P2 was used to infect Sf9 cells for protein expression and cells were harvested after 2.5 – 3 days. Cell pellets were used for protein purification or flash frozen in liquid nitrogen and stored at −80 ℃ for later use.

For constitutive expression of Flag(N)-CDK12 in HEK293 FLPN, Flag(N)-CDK12 was subcloned into pFRT vector with Gibson assembly.

For expression under Doxycycline inducible promoter, Flag(N)-CDC73, Flag(N)-CDC73ΔKIM and Flag(N)-CDC73-YY were subcloned into pFRT-TO (Addgene, #106348) vector with Gibson assembly and silent mutations were introduced for siRNA resistance.

#### Protein purification

Cell pellets were resuspended in lysis buffer: 20 mM HEPES-NaOH (pH = 8.0), 300 mM NaCl, 10 % glycerol, 1x Protease Inhibitor Mix (Roche -EDTA), 2 mM *β*-mercaptoethanol and lysed by sonication (30 pulses, 1 second each). Lysates were centrifuged at 15000 rpm for 10 min at 4 ℃ and pellets were discarded. Clarified lysates were then incubated with appropriate beads (M2 beads for Flag, Streptactin for Strep and Ni^2+^-NTA agarose for His tag purification) for 2 hours at 4 ℃ rotating. Beads were collected by centrifugation and washed with lysis buffer. Proteins were eluted with lysis buffer complemented with 0,5 mg/ml Flag peptide, 10 mM Desthiobiotin or 250 mM Imidazole according to the tag: Flag, Strep(II) or His, respectively. Elutions were further cleaned in the HiTrap-Desalt column (Cytiva) to remove elution components and reduce the NaCl concentration to 50 mM. In the case of CDK12/Cyclin K complex, a further step of ionic exchange on the HiResQ column (Cytiva) was performed before desalting. Final eluates were aliquoted, and flash frozen in liquid nitrogen before storage at – 80 ℃.

RNAPII was purified from pig thymus as previously described (*3*). The elution of the 8WG16 conjugated beads was dialyzed into 20 mM HEPES-NaOH (pH = 8.0), 150 mM NaCl, 10 % Glycerol, 2 mM *β*-mercaptoethanol and concentrated with Amicon Ultra 4 centrifugal filter (50 kDa) (Millipore). Final samples were aliquoted, and flash frozen in liquid nitrogen before storage at – 80 ℃.

#### In vitro kinase assay

Unless otherwise indicated, proteins were first dephosphorylated before performing the kinase assay. Proteins were mixed to a final concentration of 500 nM (for Coomassie stain) or 100 nM (for Western blot) in reaction buffer: 20 mM HEPES-NaOH (pH = 8.0), 50 mM NaCl, 10 % Glycerol. λPP (NEB) was added and the phosphatase reaction started by adding 4 mM MgCl_2_. Reaction was incubated for 30 min at 30 ℃. Phosphatase reaction was stopped by adding PhosSTOP 1x (Roche). Where indicated kinase inhibitors were added: 2,5 μM final concentration NVP-2 or THZ531. Kinase reaction was started by adding 1 mM ATP and incubated for the indicated times at 30 ℃. Kinase reaction was stopped by adding 10 mM EDTA. 5X SDS Loading dye (375 mM Tris 6.8, 10 % SDS, 50 % glycerol and 0.06% BPB) and 10 X reducing agent (1 M DTT) were added to the sample, boiled for 5 min at 95 ℃ before loading the gel. Products were then run in 3-8 % Tris-Acetate SDS-PAGE Gels (NuPAGE, Invitrogen) to maximize band shifts.

#### GST pulldown and NaCl gradient wash

GST-CTD was mixed with Glutathione Sepharose (Cytiva) to a final concentration of 2 µM in IP buffer: 20 mM HEPES-NaOH (pH = 8.0), 50 mM NaCl, 10 % Glycerol, 0,05 % NP-40, 2 mM *β*-mercaptoethanol, for 1 h at 4 ℃. Then CDK12 complexes were added to a final concentration of 2 µM and incubated for 1 h at 4 ℃. Input was taken and then beads were pelleted at 900 rpm for 2 min. Flow-through was collected from supernatant. Beads were washed with IP buffer with 50 mM NaCl, and the process was repeated with increasing amounts of NaCl: 100, 150, 200, 250 and 300 mM. Beads were collected at each step.

#### ELISA

96-well plates coated with NeutrAvidin (ThermoFisher; #15129) were washed 3 times with 200 µl of ELISA Washing Buffer (0.1% BSA and 0.05% Tween-20 diluted in TBS) then incubated for 2 hours at room temperature with 10 µg of the corresponding CTD peptides. Each peptide was added to the first well and then diluted 2-fold across the column for a total of 8 dilutions. After peptide incubation, plate was washed 3x times and incubated for 30 minutes at RT with primary antibodies diluted 1:2000 in 100 µl of Wash Buffer: anti-Ser2 (Active Motif; Clone 3E10), anti-Ser5 (Active Motif; Clone 3E8) and anti-Ser2+Ser5 (Cell Signaling Technologies, clone D1G3K). Plates were then washed 3x and incubated with Secondary Antibodies diluted 1:10,000 in 100 µl of Wash Buffer: anti-Rb and anti-Rat both conjugated with HRP. After 30 minutes incubation at RT, plates were washed 3x times to remove secondary antibody. For substrate developing, 100 µl of 1-Step™ TMB ELISA Substrate (Thermo; 34028) was added to each well and incubated for 5-10 mins until the desired color intensity was reached and then developing was stopped with 100 µl of Stop Solution for TMB Substrates (Thermo; N600). Plates were scanned at 450 nM using SPECTROstar Nano Microplate Reader. For each antibody treatment, samples were normalized using the min-max normalization method (X_sample_-X_min_)/(X_max_-X_min_).

#### CTD peptides

Peptides were synthesized in-house at the Francis Crick Institute and purified via HPLC. Each CTD peptide contains a biotin moiety fused to a e-aminohexanoic acid spacer (eahx) followed by 3 x tandem repeats of the aminoacid sequence YSPTSPS. Depending on the desired final peptide sequence, different permutations of phosphorylated-Serine was used for peptide synthesis. As a negative control, a scrambled CTD peptide with two phosphorylated serines per heptad was used (See Peptide Table for sequence). Peptides were resuspended in 100% DMSO and stored at −80 ℃ at a stock concentration of 10 mg/ml.

#### Peptide Table

| Name | Sequence |
| --- | --- |
| Scrambled Double Serine | Bio-eahx-PP(pS)S(pS)TYPP(pS)S(pS)TYPP(pS)S(pS)TY-COOH |
| Ser2 | Bio-eahx-Y(pS)PTSPSY(pS)PTSPSY(pS)PTSPS-COOH |
| Ser5 | Bio-eahx-YSPT(pS)PSYSPT(pS)PSYSPT(pS)PS-COOH |
| Ser2+Ser5 | Bio-eahx-Y(pS)PT(pS)PSY(pS)PT(pS)PSY(pS)PT(pS)PS-COOH |

Bio = biotin
eahx = $\epsilon$ -aminohexanoic acid

#### In vitro binding assay

Proteins were mixed to a final concentration of 1,5 µM (Strep-CTR9 alone or in PAF1 complex) and 2 µM (CDK12 complexes) in binding buffer: 20 mM HEPES-NaOH (pH = 8.0), 150 mM NaCl, 10 % Glycerol, 0,05 % NP-40, 2 mM *β*-mercaptoethanol. Input was taken and rest of the mix was incubated at 30 ℃ for 30 min and then put on ice. Equal volume of pre-washed 50 % Pierce Strep magnetic beads (Thermo, #88817) was added to the mix and incubated on ice for 2 h. Beads were washed with binding buffer complemented with 300 mM NaCl and eluted with 15 mM desthiobiotin (IBA) on ice for 1 h. Input and IP eluate was run on an SDS-PAGE gel and analyzed through Western blot.

#### Cell lines and culture

MRC5A and Flp-In T-REx HEK293 cells (R78007, Thermo Fisher Scientific, human embryonic kidney epithelial, female origin) were cultured in DMEM (1X) + GlutaMAX^TM^ (#31966-021, Gibco) supplemented with 10% v/v FBS (Tetracycline free, #S181T-500, Biowest), 100 U/mL penicillin, 100 μg/mL streptomycin at 37^◦^C and 5 % carbon dioxide.

#### Cell line generation

Flp-IN T-Rex HEK293 cell lines for Doxycycline inducible expression of Flag-CDC73 were generated as previously described (*4*). Briefly, pFRT-TO plasmid containing Flag-CDC73 was transfected with pOG44 plasmid in a 9:1 ratio using Lipofectamine 3000 (Invitrogen) according to manufacturer’s instructions. Cells were selected with 100 µg/ml hygromycin 48 h after transfection. Selected pool of cells was checked for expression of Flag-CDC73 after 24 h induction with 1 µg/ml doxycycline through Western blot.

#### siRNA knockdown

Flp-IN T-Rex HEK293 cells were seeded at a density of 2,5×10^5^ cells in a 6-well plate. The next day 2 µl of 20 µM siControl or siCDC73 (Dharmacon, J-015184-06-0005 and J-015184-08-0005) were transfected with Lipofectamine RNAiMAX (Invitrogen) according to manufacturer’s instructions. Transfection was repeated 24 h later and cells were split for the experiment the next day.

#### Whole cell extract

Cells were scraped off and centrifuged at 1000 rpm for 5 minutes. Cell pellets were resuspended and incubated in equal volume of Benzonase buffer (2 mM MgCl2, 20 mM Tris (pH 8.0), 10% glycerol, 1% Triton X-100 and 12.5 units/ml benzonase) on ice for 10 minutes. Cells were then lysed by the addition of an equal volume of 2% SDS to reach a final concentration of 1%. Samples were heated at 70 ℃ for 2 minutes. The protein concentration was determined by Bradford assay (Bio-Rad Life Science). 5X SDS Loading dye (375 mM Tris 6.8, 10 % SDS, 50 % glycerol and 0.06% BPB) and 10 X reducing agent (1 M DTT) were added to the sample, boiled for 5 min at 95 ℃ before loading the gel.

#### Chromatin fractionation

Cells were scraped off and centrifuged at 1000 rpm for 5 minutes. Cell pellets were permeabilized with CSK buffer containing 200 mM NaCl, 10 mM PIPES, 300 mM Sucrose, 1 mM MgCl2, 1 mM EDTA and 0.5% Triton X-100 on ice for 10 min. CSK fraction (supernatant) and nuclear pellet were separated by centrifugation at 900 g at 4 C for 10 min. This step can be repeated for cleaner chromatin fractions. Nuclear pellets were resuspended and incubated in equal volume of Benzonase buffer (2 mM MgCl2, 20 mM Tris (pH 8.0), 10% glycerol, 1% Triton X-100 and 12.5 units/ml benzonase) on ice for 10 minutes. The cells were then lysed by the addition of an equal volume of 2% SDS to reach a final concentration of 1%. Samples were heated at 70 ℃ for 2 minutes. The protein concentration was determined by Bradford assay (Bio-Rad Life Science). 5X SDS Loading dye (375 mM Tris 6.8, 10 % SDS, 50 % glycerol and 0.06% BPB) and 10 X reducing agent (1 M DTT) were added to the sample, boiled for 5 min at 95 ℃ before loading the gel.

#### Flag immunoprecipitation from HEK293 cells

Cell pellets were resuspended in lysis buffer: 20 mM HEPES-NaOH (pH = 8.0), 150 mM NaCl, 10 % glycerol, 1x Protease Inhibitor Mix (Roche -EDTA), 2 mM *β*-mercaptoethanol and lysed by sonication (59 pulses, 1 second each). Lysates were centrifuged at 15000 rpm for 10 min at 4 ℃ and pellets were discarded. Clarified lysates were then incubated with appropriate M2 agarose beads (#A2220, Millipore) for 2,5 hours at 4 ℃ rotating. Beads were collected by centrifugation and washed with lysis buffer. Proteins were eluted with lysis buffer complemented with 0,5 mg/ml Flag peptide for 1,5 hours at 4 ℃. 5X SDS Loading dye (375 mM Tris 6.8, 10 % SDS, 50 % glycerol and 0.06% BPB) and 10 X reducing agent (1 M DTT) were added to the sample, boiled for 5 min at 95 ℃ before loading the gel.

#### Western Blot

Proteins were transferred to nitrocellulose membranes (0,45 µm, Amersham Protran) and blocked with 5% (w/v) milk powder in TBS and 0.1% Tween 20 for 2 hours at room temperature. The membranes were incubated with antibodies against Ser2 (3E8), Ser5 (3E10) (1:1000 dilution, gift from D. Eick), Ser2 + Ser5 di-phosphorylated CTD (D1G3K, Cell Signaling), RPB1 N-terminal domain (D8L4Y, Cell Signaling), CDK9 (C12F7, Cell Signaling), Cyclin K (A301-939A, Bethyl), Lamin A (ab26300, Abcam), FLAG (D6W5B, Cell Signaling), CDK12 (A301-679A, CrkRS, Bethyl), CTR9 (A301-395A, Bethyl), PAF1 (A300-172A, Bethyl), alpha Tubulin (ab52866, Abcam) and CDC73 (A300-170A, Parafibromin, Bethyl) (1:1,000) overnight at 4 °C. Antibodies were diluted in 5 % (w/v) milk in TBS with 0.1% Tween 20 and 0.02 % Na Azide. Membranes were washed three times with TBS with 0.1% Tween 20. HRP-conjugated anti-rat secondary antibody (1: 10,000) (Sigma-Aldrich, A9037) was incubated with the membranes treated with Ser2 and Ser5 antibodies. Others were treated with HRP-conjugated anti-mouse (1: 2,000; Jackson, RRID: AB_2340770) or anti-rabbit antibody (1:20,000) (Jackson, RRID: AB_10015282) in PBS with 0.1% Tween 20 for 1 h at room temperature. SuperSignal West Pico PLUS Chemiluminescent Substrate (ThermoFisher) or, when indicated, Radiance plus (Azure biosystems) was used for detection.

#### Cell proliferation assay

Flp-In T-Rex HEK293 cells were seeded at 15×10^3^ cells in a 48-well plate (Corning, Costar #3548) in technical triplicate and biological duplicate according to condition or treatment. 1 µg/ml doxycycline was added and 24 h later the plate was placed in the Incucyte S3 (Sartorius) for live-cell imaging. Pictures were taken every 4 h for 5 days and growth curves were plotted with resulting confluence measurements.

#### TTchem-seq

Protocol is adapted from (*5*). Flp-In T-Rex HEK293 cells were subjected to switchover in a 150 mm plate (Corning) as previously described. Nascent RNA was then labelled using a 15-minute pulse of 4-thiouridine (4sU; Glentham Life Sciences, #GN6085), after which the reaction was stopped by aspiration of media and addition of 1mL Trizol (Thermo Fisher Scientific, #15596026). For total RNA isolation, one-fifth of chloroform (Alfa Aesar; Thermo Fisher Scientific, #43685) was added to lysates and shaken well, followed by addition of the upper aqueous phase and an equal volume of chloroform/isoamyl alcohol (24:1; Sigma-Aldrich, #C0549) to MaXtract high-density phase-lock tubes (Qiagen, #129046), that were previously prepared according to manufacturer’s instructions. Lysates were then centrifuged (12000 g, 5 minutes at 4 °C) and upper aqueous phase was transferred to a new tube and 1.1X volumes of 100 % isopropanol (Thermo Fisher Scientific, #P/7500/PC17) was added, followed by shaking for 30 seconds, incubation at room temperature for 20 minutes and centrifugation (12000 g, 20 minutes at 4 °C). Supernatant was removed and RNA pellets were washed using 750 µL 85 % (v/v) ethanol, centrifuged (7500 g, 5 minutes at 4 °C), air dried at room temperature and resuspended in 100 µL Ultrapure DNase/RNase-free distilled water. As an external reference control, *S. cerevisiae* (strain BY4741, W303, MATa, his3D1, leu2D0, met15D0, ura3D0) nascent RNA was spiked-in to each sample. Briefly, *S. cerevisiae* cells were grown to mid-log phase (OD600 of 0.5) in YPD medium, after which nascent RNA was labelled using a 6-minute pulse of 5 mM 4-thiouracil (4TU). Total RNA was then isolated using PureLink RNA Mini kit (Thermo Fisher Scientific, #12183020) according to the manufacturer’s instruction for the enzymatic protocol. 500 ng of *S. cerevisiae* total RNA was spiked-in to 100 µg of *H. sapiens* total RNA, material was adjusted to 100µL and fragmented by the addition of 20 µL 1 M NaOH (Sigma-Aldrich, #72068) and incubation on ice for 20 minutes. Fragmentation was stopped by the addition of 80 µL 1 M Tris HCl pH 6.8 and material was purified twice using the Micro Bio-Spin P-30 Gel Columns (BioRad, #7326223) according to manufacturer’s instructions. Thiol-specific biotinylation of RNA was performed using 10 mM Tris-HCl pH 7.4, 1 mM EDTA and 5 mg MTSEA biotin-XX linker (Biotium, #BT90066) in a final volume of 250 µL and incubation at room temperature for 30 minutes in the dark. Biotinylated RNA was then purified by addition of 250μL of phenol/chloroform/isoamyl alcohol (25:24:1 (vol/vol); Thermo Fisher, #15593031), followed by shaking and then addition to MaXtract high-density phase-lock tubes (Qiagen, #129046) that were previously prepared according to manufacturer’s instructions.

Upper aqueous phase was transferred to a new tube and biotinylated RNA was precipitated using one-tenth the volume 5 M NaCl and 1.1 X volume 100 % isopropanol, followed by incubation room temperature for 10 minutes and centrifugation (20000 g, 20 minutes at 4 °C). Supernatant was discarded and RNA pellet was washed using 500 µL 85 % (v/v) ethanol, centrifuged (20000 g, 5 minutes at 4 °C), air dried at room temperature, resuspended in 50 µL Ultrapure DNase/RNase-free distilled water and denatured by incubating for 10 minutes at 65 °C in the dark. Biotinylated RNA was separated from the total RNA pool by addition of 200 μl μMACS Streptavidin MicroBeads (Miltenyi, 130-074-101) and incubation for 15 minutes at room temperature with constant rotation. Beads were then applied to a μColumn pre-equilibrated with 100 µL nucleic acid equilibration buffer and immobilised on a magnetic stand, washed twice with pre-warmed (55 °C) wash buffer (100 mM Tris HCl pH 7.4 10 mM EDTA, 1 M NaCl and 0.1 % (v/v) Tween 20) and biotinylated RNA was eluted by two additions of 100 μl 100 mM DTT 5 minutes apart. Purification of biotinylated RNA was performed using the RNeasy MinElute kit (QIAGEN, 74204) according to manufacturer’s instructions, where 1050 μl 100% ethanol and 750 μl RLT buffer was adder per 20 μl reaction to precipitate RNA < 200 nt. Biotinylated RNA quantity and fragment size was assessed with the Agilent TapeStation 4150 and Qubit 3.0 Fluorometer respectively. Sequencing libraries were prepared from biotinylated RNA using the NEBNext Ultra II Directional RNA Library Prep Kit for Illumina (without additional fragmentation) (NEB, # E7760S) and NEBNext Multiplex Oligos for Illumina (Unique Dual Index UMI adapters RNA Set 1) (NEB, #E7416S) and sequenced on the NextSeq 2000 (P3 50 kit).

#### TT_chem_-seq analysis

Paired-end reads were demultiplexed using bcl2fastq (v2.20.0.422) and resultant FASTQ files were quality checked using MultiQC (v1.11). Unique molecular identifiers (UMIs) present in R2 were appended to R1, and resulting FASTQ file was mapped to GRCh38/hg38 and sacCer3 genomes using hisat2 (v2.2.1) with standard single-end read settings. Mapped reads in SAM format were converted to BAM (view), which were then sorted (sort) and indexed (index) using samtools (v1.20) with standard settings, and PCR duplicates were removed using UMI tools (v1.1.4) deduplication function (dedup). For external reference normalisation using yeast spike-in material, reads mapping to hg38 and sacCer3 chromosomes were counted using FeatureCounts from the subread package (v2.0.6; featureCounts -O -M -T 32 -s 0 -F SAF), after which edgeR (v3.40.2) on Rstudio (v4.2.1) was used to determine scaling factors for BAM to bigWig file conversion using the bamCoverage function from the deeptools package (v3.5.5, bamCoverage -of bigwig -bs 50 -p max --normalizeUsing CPM --smoothLength 150 -- filterRNAstrand forward/reverse –scaleFactor X). Metagene analysis across gene intervals using scaled bigWig files was performed using the computeMatrix function (scale-regions -- regionBodyLength 5000 --upstream 2000 --downstream 2000 --binSize 50 -p 32 – missingDataAsZero), followed by subsetting of files according to stand (computeMatrixOperations subset), and direction of transcription (computeMatrixOperations filterStrand), after which forward and reverse files were concatenated according to direction transcription (computeMatrixOperations rbind), all using the deeptools package (v3.5.5). Matrix files were then visualised using ggplot2 (v3.5.1) on R (v4.2.1).

---

**Fig. S1.**
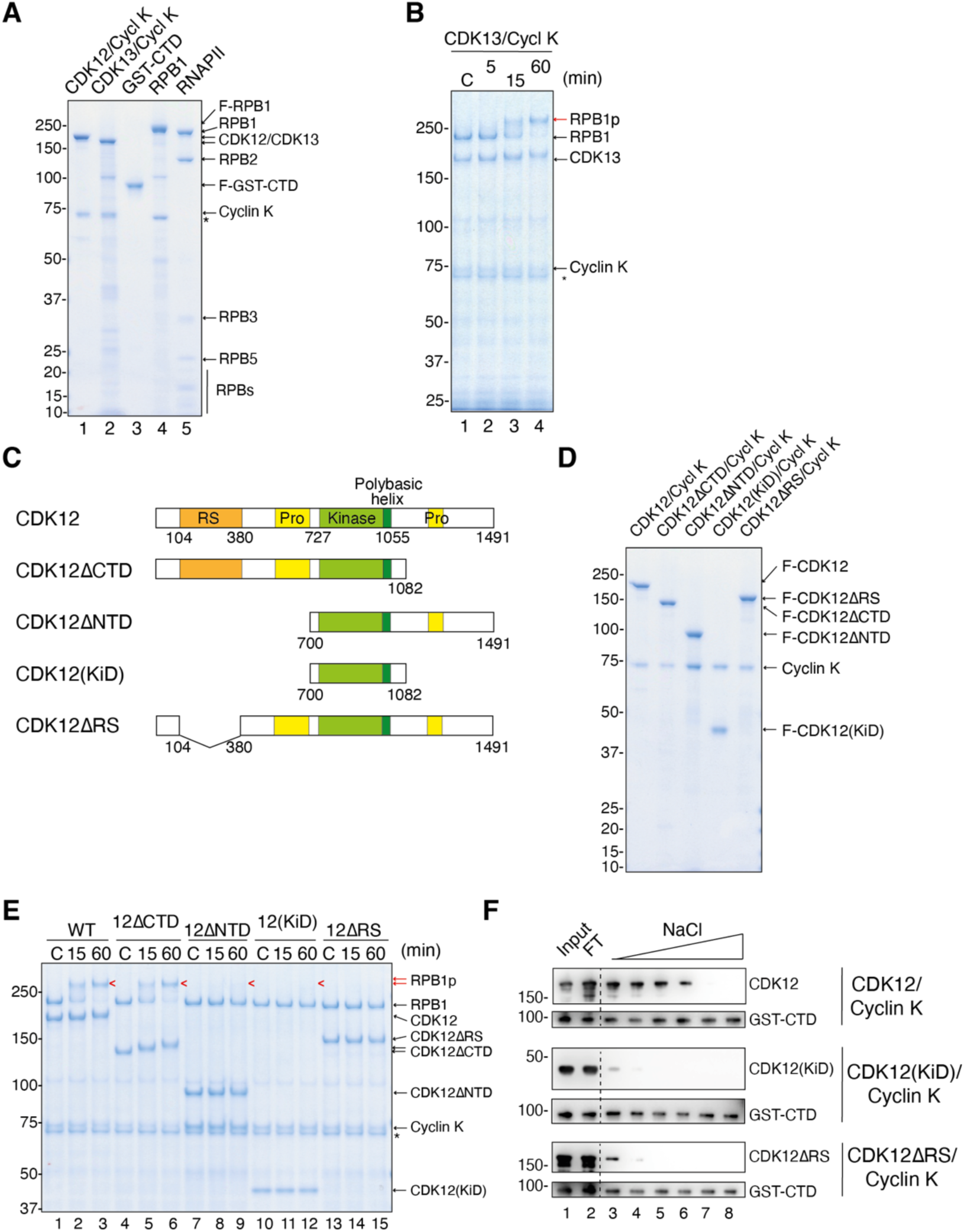
**>Related to Figure 1.** (**A**) Coomassie blue stained gel showing purified CDK12/Cyclin K, CDK13/Cyclin K, GST-CTD, Flag-RPB1 and RNAPII (from pig thymus). (**B**) Coomassie blue stained SDS-PAGE of an *in vitro* kinase assay using CDK13/Cyclin K and full-length RPB1 for the indicated times. C: control with no ATP added. (**C**) Schematic representation of the domain organization of wild type and truncated constructs used in this study. CTD: C-terminal domain, NTD: N-terminal domain, KiD: kinase domain and RS: arginine and serine-rich domain. (**D**) Coomassie blue stained gel showing purified CDK12/Cyclin K WT or truncated mutant forms as in (C). (**E**) Coomassie blue stained SDS-PAGE of an *in vitro* kinase assay using CDK12/Cyclin K wild-type (WT) or truncated forms as in (C) on full-length RPB1 for the indicated times in minutes (min). C: control with no ATP added. (**F**) Western blot of in vitro binding reaction, mixing GST-CTD with the indicated forms of CDK12/Cyclin K, followed by binding to glutathione sepharose, and step-wise washes with increasing NaCl concentrations. FT: flow-through.

**Fig. S2.**
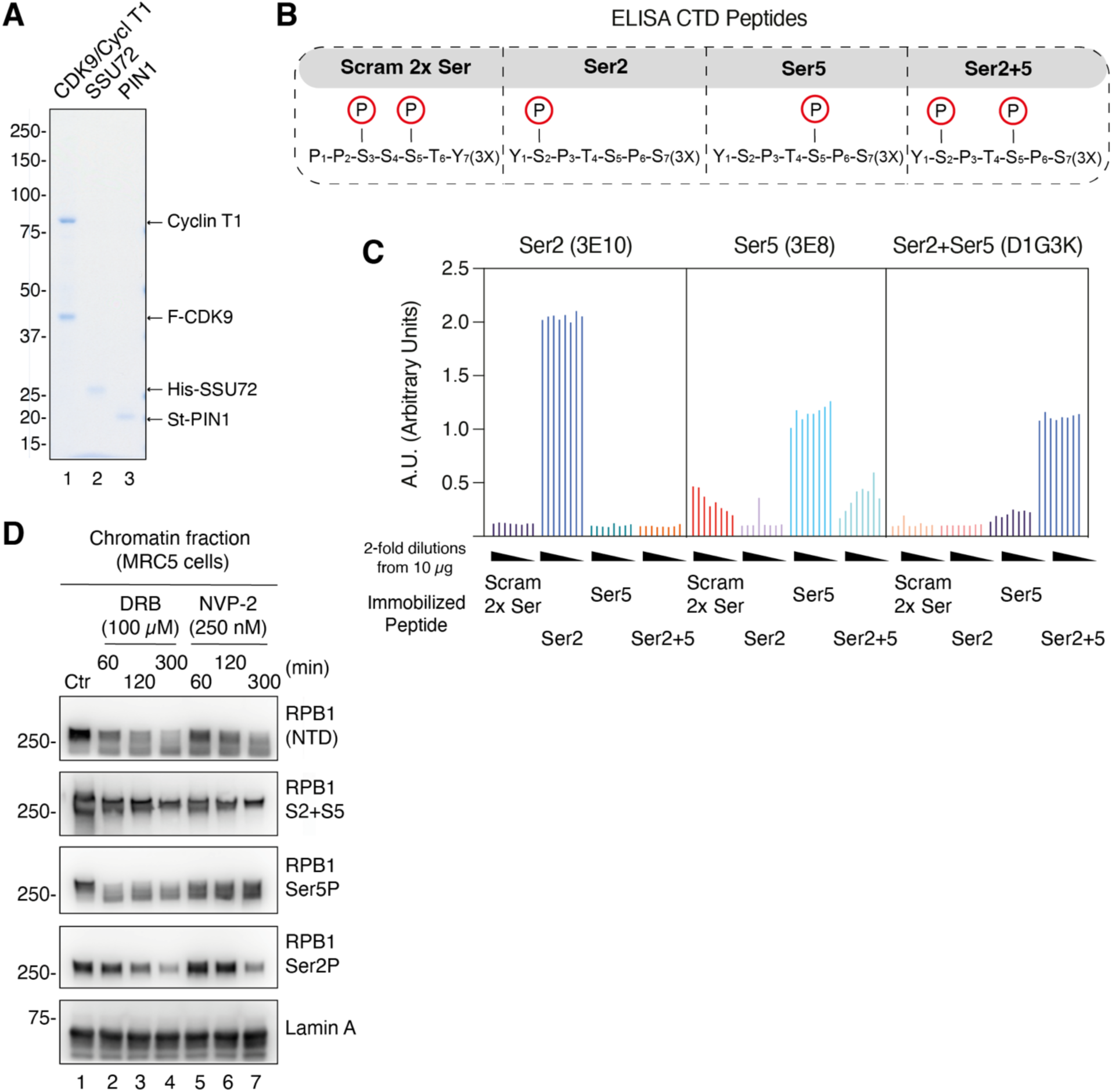
**Related to Figure 1.** (**A**) Coomassie blue stained gel showing purified CDK9/Cyclin T1, His-SSU72 and Strep(II)-PIN1. (**B**) Schematic representation of the chemically synthesized phosphorylated CTD (3x) peptides used for the ELISA. (**C**) ELISA showing the binding affinities of phosphor-CTD specific antibodies used in this study: anti-Ser2 (Clone 3E10), anti-Ser5 (Clone 3E8) and anti-Ser2+Ser5 (clone D1G3K) and chemically synthesized CTD peptides containing Ser5P, Ser2P, Ser2P-Ser5P or scrambled di-phosphorylation. **(D)** Western blot of the chromatin fractions of MRC5 cells treated with DRB (100 μM) or NVP-2 (250 nM) for the indicated times. Ctr: no drug added.

**Fig. S3.**
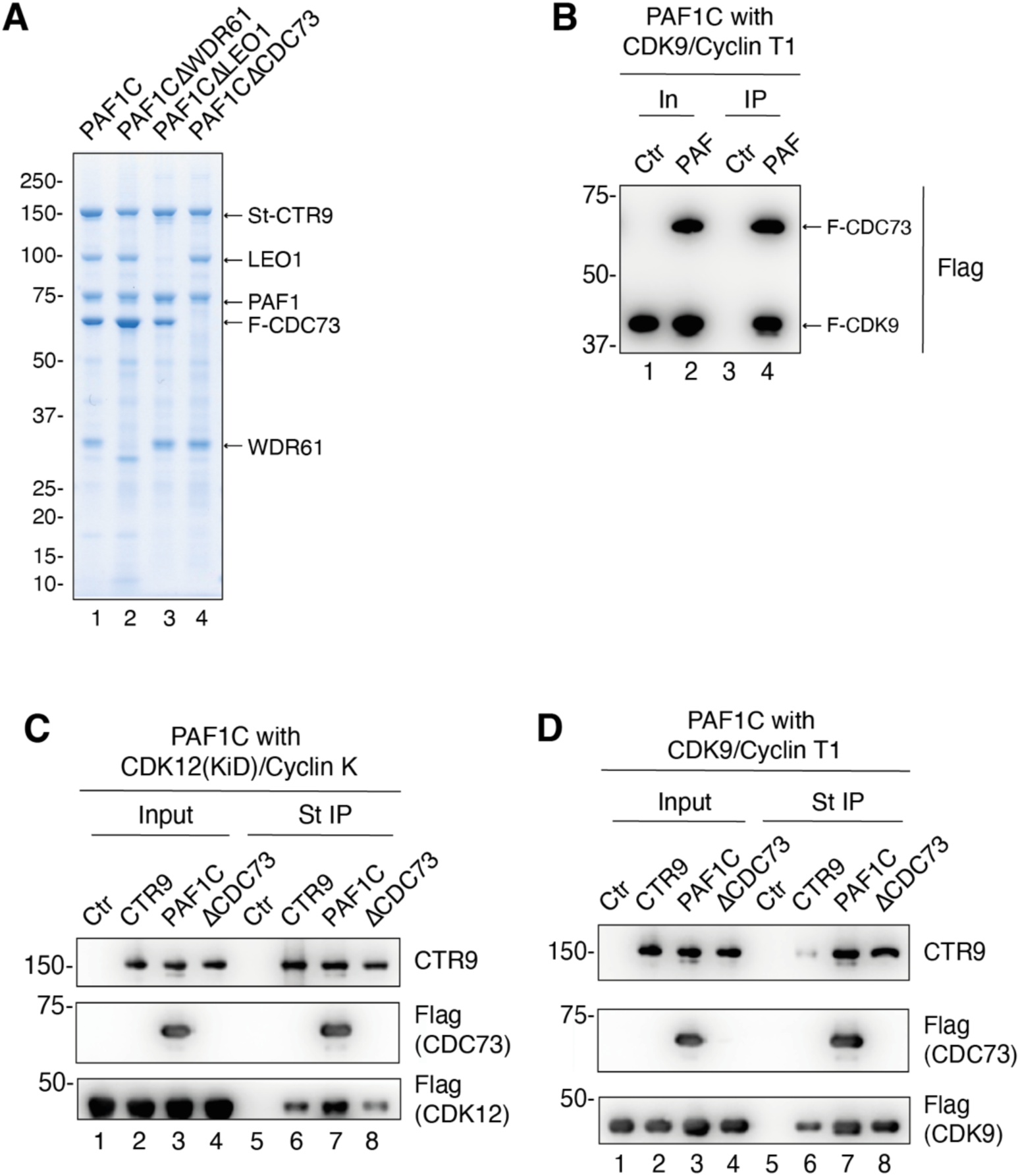
**Related to Figure 2.** (**A**) Coomassie blue stained gel showing purified PAF1 complex or incomplete PAF1C missing either WDR61, LEO1 or CDC73 subunits (**B**) Western blot of an *in vitro* binding reaction between the PAF1 complex and CDK9/Cyclin T1 followed by Strep IP. Ctr: no PAF1 complex added. (**C**) Western blot of an *in vitro* binding reaction between either St-CTR9, PAF1 complex or PAF1CΔCDC73 and CDK12(KiD)/Cyclin K complex followed by Strep IP. Ctr: no PAF1 complex added. (**D**) Western blot of an *in vitro* binding reaction between either St-CTR9, PAF1 complex or PAF1CΔCDC73 and CDK9/Cyclin T1 complex followed by Strep IP. Ctr: no PAF1 complex added.

**Fig. S4.**
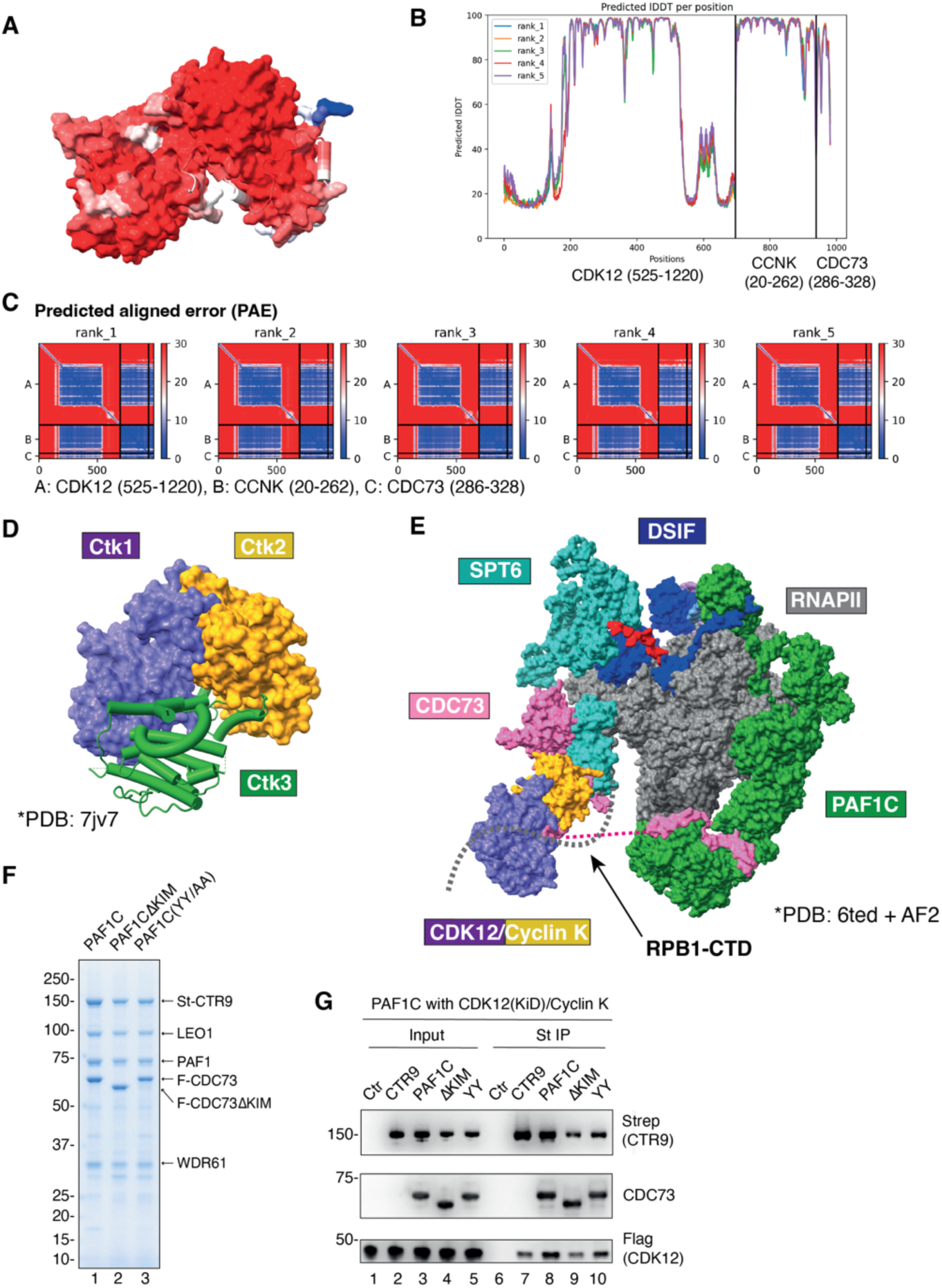
**Related to Figure 3.** (**A**) AlphaFold2 (AF2) structural model of the trimeric complex of the CDK12 kinase domain, the Cyclin K cyclin fold, and the CDC73-KIM, colored according to the prediction confidence (b-factor, red indicates high confidence). (**B**) Predicted local distance difference test (pLDDT) per position for the five AF2 models of the trimeric complex. pLDDT is > 80. (**C**) Predicted aligned error (PAE) of the five models generated by AF2. This contrasts with other contacts across subunits of the trimer, where predicted aligned error was low across different model predictions. (**D**) Crystal structure model of the yeast CTDK-1 trimer, showing Ctk1 (purple), Ctk2 (yellow) and Ctk3 (green) (PDB accession code: 7JV7). (**E**) Cryo-EM model of the RNAPII elongation complex containing DSIF (dark blue), SPT6 (light blue) and the PAF1 complex (green) (PDB accession code: 6TED). The CDC73 C-terminus (pink) was modelled onto the SPT6 C-terminal SH2 domain using AF2 and superimposed. CDK12/Cyclin K/CDC73-KIM was then added to the model in an approximate location in the vicinity of RNAPII CTD. (**F**) Coomassie blue stained gel showing purified PAF1 complex, PAF1CΔKIM and PAF1C-YY/AA. (**G**) Western blot of an *in vitro* binding reaction between either St-CTR9, PAF1 complex, PAF1CΔKIM or PAF1C-YY/AA and CDK12(KiD)/Cyclin K complex followed by Strep IP. Ctr: no PAF1 complex added.

**Fig. S5.**
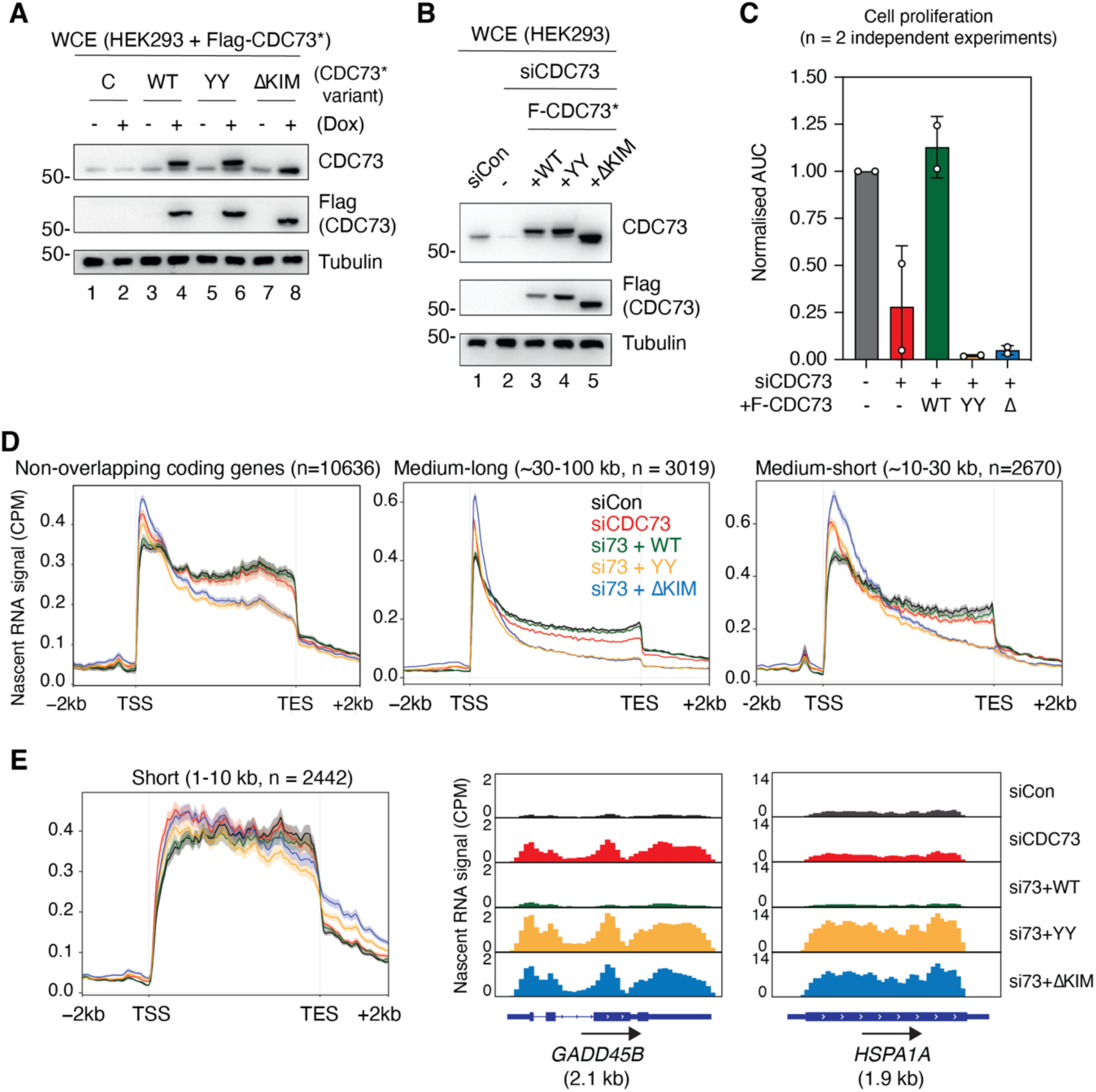
**Related to Figure 4.** (**A**) Western blot of whole cell extracts (WCE) of HEK293 cell lines with or without 1µg/mL Dox for 24 hours (h) to show the expression of recombinant Flag-CDC73 siRNA resistant (Flag-CDC73*) WT and CDC73-KIM mutant forms (**B**) Western blot validation of the switchover system used for the proliferation assays (**C**) Quantification of the proliferation assays performed with the switchover system of CDC73. Area under the curve (AUC) showed for each cell line/treatments in n = 2 independent experiments. (**D**) (Left) Metagene analysis of non-overlapping coding gene intervals (n = 10636) using library and spike-in normalized nascent RNA read counts obtained from transient transcriptome chem sequencing (TT_chem_-seq). Line indicates mean signal of experiment performed in technical triplicate. Shaded area indicates standard error. Si73: siRNA targeting CDC73, CPM: counts per million, TSS: transcription start site, TES: transcription end site, kb: kilobases. (Middle and Right) Metagene analysis of non-overlapping coding gene intervals of medium-long genes (n = 3019) (Middle), and medium-short (n = 2670) (Right) genes. CPM: counts per million, TSS: transcription start site, TES: transcription end site, kb: kilobases. (**E**) (Left) As in (D) but for short genes (n = 2442). (Right) Representative images of library- and spike-in normalized nascent RNA read counts across *GADD45B* and *HSPA1A*. Direction of transcription indicated by arrow.

## Notes

### Competing Interest Statement

The authors have declared no competing interest.

## References and Notes

1. R. G. Roeder, 50+ years of eukaryotic transcription: an expanding universe of factors and mechanisms. Nat Struct Mol Biol 26, 783–791 (2019).

2. X. Liu, D. A. Bushnell, R. D. Kornberg, RNA polymerase II transcription: structure and mechanism. Biochim Biophys Acta 1829, 2–8 (2013).

3. S. Osman, P. Cramer, Structural Biology of RNA Polymerase II Transcription: 20 Years On. Annu Rev Cell Dev Biol, (2020).

4. J. Q. Svejstrup, The RNA polymerase II transcription cycle: cycling through chromatin. Biochim Biophys Acta 1677, 64–73 (2004).

5. S. Buratowski, Progression through the RNA polymerase II CTD cycle. Mol Cell 36, 541–546 (2009).

6. D. Blazek et al., The Cyclin K/Cdk12 complex maintains genomic stability via regulation of expression of DNA damage response genes. Genes Dev 25, 2158–2172 (2011).

7. B. Bartkowiak et al., CDK12 is a transcription elongation-associated CTD kinase, the metazoan ortholog of yeast Ctk1. Genes Dev 24, 2303–2316 (2010).

8. P. K. Parua, R. P. Fisher, Dissecting the Pol II transcription cycle and derailing cancer with CDK inhibitors. Nat Chem Biol 16, 716–724 (2020).

9. S. Basu, J. Greenwood, A. W. Jones, P. Nurse, Core control principles of the eukaryotic cell cycle. Nature 607, 381–386 (2022).

10. R. Abdella et al., Structure of the human Mediator-bound transcription preinitiation complex. Science 372, 52–56 (2021).

11. Y. J. Kim, S. Bjorklund, Y. Li, M. H. Sayre, R. D. Kornberg, A multiprotein mediator of transcriptional activation and its interaction with the C-terminal repeat domain of RNA polymerase II. Cell 77, 599–608 (1994).

12. P. J. Robinson et al., Structure of a Complete Mediator-RNA Polymerase II Pre-Initiation Complex. Cell 166, 1411–1422 e1416 (2016).

13. M. T. M. Sogaard, J. Q. Svejstrup, Hyperphosphorylation of the C-terminal Repeat Domain of RNA Polymerase II Facilitates Dissociation of Its Complex with Mediator. J Biol Chem 282, 14113–14120 (2007).

14. G. Garg et al., Structural insights into human co-transcriptional capping. Mol Cell 83, 2464–2477 e2465 (2023).

15. E. J. Cho, T. Takagi, C. R. Moore, S. Buratowski, mRNA capping enzyme is recruited to the transcription complex by phosphorylation of the RNA polymerase II carboxy-terminal domain [see comments]. Genes Dev 11, 3319–3326 (1997).

16. S. McCracken et al., 5’-Capping enzymes are targeted to pre-mRNA by binding to the phosphorylated carboxy-terminal domain of RNA polymerase II. Genes Dev 11, 3306–3318 (1997).

17. L. J. Core et al., Defining the status of RNA polymerase at promoters. Cell Rep 2, 1025–1035 (2012).

18. M. DeBerardine, G. T. Booth, P. P. Versluis, J. T. Lis, The NELF pausing checkpoint mediates the functional divergence of Cdk9. Nat Commun 14, 2762 (2023).

19. C. W. Bacon, I. D’Orso, CDK9: a signaling hub for transcriptional control. Transcription 10, 57–75 (2019).

20. S. M. Vos, L. Farnung, H. Urlaub, P. Cramer, Structure of paused transcription complex Pol II-DSIF-NELF. Nature 560, 601–606 (2018).

21. S. M. Vos et al., Structure of activated transcription complex Pol II-DSIF-PAF-SPT6. Nature 560, 607–612 (2018).

22. A. D. Wier, M. K. Mayekar, A. Heroux, K. M. Arndt, A. P. VanDemark, Structural basis for Spt5-mediated recruitment of the Paf1 complex to chromatin. Proc Natl Acad Sci U S A 110, 17290–17295 (2013).

23. A. M. Francette, S. A. Tripplehorn, K. M. Arndt, The Paf1 Complex: A Keystone of Nuclear Regulation Operating at the Interface of Transcription and Chromatin. J Mol Biol 433, 166979 (2021).

24. M. Yu et al., RNA polymerase II-associated factor 1 regulates the release and phosphorylation of paused RNA polymerase II. Science 350, 1383–1386 (2015).

25. M. Tellier et al., CDK12 globally stimulates RNA polymerase II transcription elongation and carboxyl-terminal domain phosphorylation. Nucleic Acids Res 48, 7712–7727 (2020).

26. C. A. Bosken et al., The structure and substrate specificity of human Cdk12/Cyclin K. Nat Commun 5, 3505 (2014).

27. B. Bartkowiak, C. M. Yan, E. J. Soderblom, A. L. Greenleaf, CDK12 Activity-Dependent Phosphorylation Events in Human Cells. Biomolecules 9, (2019).

28. A. P. Chirackal Manavalan et al., CDK12 controls G1/S progression by regulating RNAPII processivity at core DNA replication genes. EMBO Rep 20, e47592 (2019).

29. Z. Fan et al., CDK13 cooperates with CDK12 to control global RNA polymerase II processivity. Sci Adv 6, eaaz5041 (2020).

30. Y. Liu et al., Discovery of MFH290: A Potent and Highly Selective Covalent Inhibitor for Cyclin-Dependent Kinase 12/13. J Med Chem 63, 6708–6726 (2020).

31. B. Bartkowiak, C. Yan, A. L. Greenleaf, Engineering an analog-sensitive CDK12 cell line using CRISPR/Cas. Biochim Biophys Acta 1849, 1179–1187 (2015).

32. K. Liang et al., Characterization of human cyclin-dependent kinase 12 (CDK12) and CDK13 complexes in C-terminal domain phosphorylation, gene transcription, and RNA processing. Mol Cell Biol 35, 928–938 (2015).

33. A. K. Greifenberg et al., Structural and Functional Analysis of the Cdk13/Cyclin K Complex. Cell Rep 14, 320–331 (2016).

34. M. Krajewska et al., CDK12 loss in cancer cells affects DNA damage response genes through premature cleavage and polyadenylation. Nat Commun 10, 1757 (2019).

35. R. D. Chapman et al., Transcribing RNA polymerase II is phosphorylated at CTD residue serine-7. Science 318, 1780–1782 (2007).

36. A. M. Sanchez, A. Garg, S. Shuman, B. Schwer, Genetic interactions and transcriptomics implicate fission yeast CTD prolyl isomerase Pin1 as an agent of RNA 3’ processing and transcription termination that functions via its effects on CTD phosphatase Ssu72. Nucleic Acids Res 48, 4811–4826 (2020).

37. M. Heidemann, C. Hintermair, K. Voss, D. Eick, Dynamic phosphorylation patterns of RNA polymerase II CTD during transcription. Biochim Biophys Acta 1829, 55–62 (2013).

38. M. Mirdita et al., ColabFold: making protein folding accessible to all. Nature Methods 19, 679–682 (2022).

39. R. Evans, et al., Protein complex prediction with AlphaFold-Multimer. bioRxiv, 2021.2010.2004.463034 (2022).

40. J. Jumper et al., Highly accurate protein structure prediction with AlphaFold. Nature 596, 583–589 (2021).

41. Y. Xie et al., Structure and activation mechanism of the yeast RNA Pol II CTD kinase CTDK-1 complex. Proc Natl Acad Sci U S A 118, (2021).

42. S. M. Vos, L. Farnung, A. Linden, H. Urlaub, P. Cramer, Structure of complete Pol II-DSIF-PAF-SPT6 transcription complex reveals RTF1 allosteric activation. Nat Struct Mol Biol 27, 668–677 (2020).

43. M. A. Ellison et al., Spt6 directly interacts with Cdc73 and is required for Paf1 complex occupancy at active genes in Saccharomyces cerevisiae. Nucleic Acids Res, (2023).

44. H. Ehara, T. Kujirai, M. Shirouzu, H. Kurumizaka, S. I. Sekine, Structural basis of nucleosome disassembly and reassembly by RNAPII elongation complex with FACT. Science 377, eabp9466 (2022).

45. S. J. Dubbury, P. L. Boutz, P. A. Sharp, CDK12 regulates DNA repair genes by suppressing intronic polyadenylation. Nature 564, 141–145 (2018).

46. L. H. Gregersen, R. Mitter, J. Q. Svejstrup, Using TTchem-seq for profiling nascent transcription and measuring transcript elongation. Nat Protoc 15, 604–627 (2020).

47. B. Schwalb et al., TT-seq maps the human transient transcriptome. Science 352, 1225–1228 (2016).

48. R. Schuller et al., Heptad-Specific Phosphorylation of RNA Polymerase II CTD. Mol Cell 61, 305–314 (2016).

49. E. Vojnic, B. Simon, B. D. Strahl, M. Sattler, P. Cramer, Structure and Carboxyl-terminal Domain (CTD) Binding of the Set2 SRI Domain That Couples Histone H3 Lys36 Methylation to Transcription. Journal of Biological Chemistry 281, 13–15 (2006).

50. L. H. Gregersen et al., SCAF4 and SCAF8, mRNA Anti-Terminator Proteins. Cell 177, 1797–1813 e1718 (2019).

51. D. Lopez Martinez, J. Q. Svejstrup, Mechanisms of RNA Polymerase II termination at the 3’-end of genes. J Mol Biol, 168735 (2024).

52. L. Davidson, L. Muniz, S. West, 3’ end formation of pre-mRNA and phosphorylation of Ser2 on the RNA polymerase II CTD are reciprocally coupled in human cells. Genes Dev 28, 342–356 (2014).

53. J. M. Lee, A. L. Greenleaf, CTD kinase large subunit is encoded by CTK1, a gene required for normal growth of Saccharomyces cerevisiae. Gene Expr 1, 149–167 (1991).

54. J. E. Krebs, B. Lewin, S. T. Kilpatrick, E. S. Goldstein, Lewin’s genes XI. (Jones & Bartlett Learning, Burlington, Mass., ed. 11th, 2014), pp. xxvii, 940 p.

55. G. Fuchs et al., 4sUDRB-seq: measuring genomewide transcriptional elongation rates and initiation frequencies within cells. Genome Biol 15, R69 (2014).

56. C. G. Danko et al., Signaling pathways differentially affect RNA polymerase II initiation, pausing, and elongation rate in cells. Mol Cell 50, 212–222 (2013).

57. I. Jonkers, H. Kwak, J. T. Lis, Genome-wide dynamics of Pol II elongation and its interplay with promoter proximal pausing, chromatin, and exons. Elife 3, e02407 (2014).

58. M. Saponaro et al., RECQL5 Controls Transcript Elongation and Suppresses Genome Instability Associated with Transcription Stress. Cell 157, 1037–1049 (2014).

59. R. Düster et al., Structural basis of Cdk7 activation by dual T-loop phosphorylation. Nature Communications 15, (2024).

60. S. Peissert, A. Schlosser, R. Kendel, J. Kuper, C. Kisker, Structural basis for CDK7 activation by MAT1 and Cyclin H. Proc Natl Acad Sci U S A 117, 26739–26748 (2020).

61. T. H. Tahirov et al., Crystal structure of HIV-1 Tat complexed with human P-TEFb. Nature 465, 747–751 (2010).

62. U. Schulze-Gahmen et al., The AFF4 scaffold binds human P-TEFb adjacent to HIV Tat. Elife 2, e00327 (2013).

## References

1. F. Weissmann, J. M. Peters, Expressing Multi-subunit Complexes Using biGBac. Methods Mol Biol 1764, 329–343 (2018).

2. F. Weissmann et al., biGBac enables rapid gene assembly for the expression of large multisubunit protein complexes. Proc Natl Acad Sci U S A 113, E2564–2569 (2016).

3. X. Hu et al., A Mediator-responsive form of metazoan RNA polymerase II. Proc Natl Acad Sci U S A 103, 9506–9511 (2006).

4. L. H. Gregersen et al., SCAF4 and SCAF8, mRNA Anti-Terminator Proteins. Cell 177, 1797–1813 e1718 (2019).

5. L. H. Gregersen, R. Mitter, J. Q. Svejstrup, Using TTchem-seq for profiling nascent transcription and measuring transcript elongation. Nat Protoc 15, 604–627 (2020).

